# A Long-Circulating Vector for Aptamers Based upon Polyphosphodiester-Backboned Molecular Brushes

**DOI:** 10.1101/2022.06.30.498214

**Authors:** Yuyan Wang, Dali Wang, Jiachen Lin, Zidi Lyu, Peiru Chen, Tingyu Sun, Chenyang Xue, Mehrnaz Mojtabavi, Armin Vedadghavami, Zheyu Zhang, Ruimeng Wang, Lei Zhang, Christopher Park, Gyu Seong Heo, Yongjian Liu, Sijia Dong, Ke Zhang

## Abstract

Aptamers face challenges for use outside the ideal conditions in which they are developed. These difficulties are most palpable *in vivo* due to nuclease activities, rapid clearance, and off-target binding. Herein, we demonstrate that a polyphosphodiester-backboned molecular brush can suppress enzymatic digestion, reduce non-specific cell uptake, enable long blood circulation, and rescue the bioactivity of a conjugated aptamer *in vivo*. The backbone along with the aptamer is assembled via solid-phase synthesis, followed by installation of poly(ethylene glycol) (PEG) side chains using a two-step process with near-quantitative efficiency. The synthesis allows for precise control over polymer size and architecture. Consisting entirely of building blocks that are generally recognized as safe for therapeutics, this novel molecular brush is expected to provide a highly translatable route for aptamer-based therapeutics.

## Introduction

Aptamers are single-stranded oligonucleotides that can fold into defined secondary structures with a high binding affinity to their targets.^[1]^ Compared to antibodies, aptamers enjoy a wide range of advantages including better production scalability, lower development cost, non-immunogenicity, less susceptibility to biological contamination, better tissue penetration, and more ease to develop an antidote.^[2]^ However, these advantages are offset by two serious drawbacks: poor *in vivo* stability and difficult pharmacological properties.^[3]^ As a result, aptamers have seen very limited commercial success, with only one drug on the market: Macugen®, a PEGylated, multi-modified aptamer that is delivered locally (intravitreal) to treat age-related macular degeneration.^[4]^ Even in this space, Macugen® is facing severe competition from antibody alternatives (Lucentis® or off-label use of Avastin®) that bind to the same target, vascular endothelial growth factor A.^[5]^

Aptamers that bind to a specific target are selected from a random sequence library by a process termed systematic evolution of ligands by exponential enrichment (SELEX).^[6]^ Advances in SELEX technologies have enabled a small number of chemical modifications such as 2’-fluoro, 2’-amino, and α-nucleoside thiotriphosphates (Sp) to be incorporated into the selection process,^[7]^ which render the resulting aptamers more resistant towards degrading enzymes. Aptamers can also be tested post-SELEX for tolerance of modifications that can further enhance their properties, such as 2’-OMe substitution of purines, 3’-capping, and bioconjugation (e.g. with lipids, cholesterol, or polymers).^[8]^ Further, aptamers have been prepared using the enantiomeric form of natural nucleic acids (Spiegelmer®), which makes such aptamers completely unsusceptible to nucleases.^[9]^ Together, these advances have considerably addressed the *in vivo* stability aspect of aptamers, leaving pharmacological limitations a primary hurdle for clinical translation.

We have recently reported that a bottlebrush polymer with dense PEG side chains can enhance the plasma pharmacokinetics (PK) and bioavailability of conjugated antisense oligonucleotides by steric inhibition of specific/non-specific nucleic acid-protein interactions.^[10]^ Such steric selectivity greatly reduces side effects associated with oligonucleotide therapeutics, including coagulopathy and unwanted activation of the immune system. In addition, the nanoscopic nature of the conjugate produces a novel biodistribution profile, elevates blood circulation times, and augments tissue retention (up to 15 weeks post intravenous injection). These improvements result in massively boosted antisense activities *in vivo*, with 1-2 orders of magnitude reduction in dosage requirement in certain cancer xenograft models. However, our initial bottlebrush polymer system cannot be readily applied to aptamers due to their tendency to undergo endocytosis, which we speculate stems from the hydrophobic polynorbornene (PN) backbone of the bottlebrush polymer (Scheme 1).^[11]^ Thus, for applications involving aptamer targeting of extracellular species, a new chemistry that retains the favorable pharmacological properties of prototypical bottlebrush-oligonucleotide conjugates but suppresses cellular uptake is preferrable. Herein, we report a novel bottlebrush polymer system with a completely hydrophilic, phosphodiester backbone capable of extended blood circulation and restoring aptamer bioactivity *in vivo*.

## Results and Discussion

The phosphodiester backbone of the bottlebrush polymer is assembled by stepwise condensation of an Fmoc-protected phosphoramidite derived from serinol (Scheme S1), a common intermediate for pharmaceuticals.^[12]^ Because the synthesis of the polymer backbone shares the same chemistry as oligonucleotide synthesis, the aptamer portion can be prepared as part of the polymer backbone, eliminating subsequent aptamer conjugation and purification steps. For proof of concept, a poly(serinol phosphodiester) (PSP) backbone of 30 repeating units with two dT_15_ strands flanking each terminus of the backbone was synthesized (dT_15_-*b*-PSP_30_-*b*-dT_15_). Following the synthesis, the Fmoc groups protecting the serinol amines were removed, and the strand was purified by reversed-phase high performance liquid chromatography (RP-HPLC, Figure S1).

To construct the bottlebrush segment, the amine groups were derivatized with *N*-hydroxysuccinimide (NHS)-terminated PEG in a two-stage process. In the first stage, the purified backbone was treated with one equiv. of 10 kDa PEG succinimidyl glutaramide in 1× phosphate buffered saline (PBS, pH 7.4) at 4 °C overnight. Next, the product from the first stage was lyophilized and reacted with another equiv. of PEG in anhydrous *N,N*-dimethylformamide (DMF) for 60 h at room temperature. The two-stage PEGylation is necessary because NHS esters hydrolyze in an aqueous buffer leading to unsatisfactory coupling efficiencies, but without the initial aqueous coupling step, the anionic backbone/oligonucleotide sequence is insoluble in DMF. With the two-step process, near-quantitative coupling yields (∼95% conversion of serinol amines) were obtained as determined by a 2,4,6-trinitrobenzene sulfonic acid (TNBSA) assay using glycine as a standard (Figure S2 and Table S2). The incremental PEGylation produced incremental increases in molecular weight (MW) after each coupling reaction, as observed by aqueous gel permeation chromatography (GPC) (Figure 1a and 1b). Atomic Force Microscopy (AFM) measurement of the final structure, dT_15_-*b*-(PSP_30_-*g*-PEG)-*b*-dT_15_, shows a spherical morphology with a dry-state diameter of 21±3 nm (Figure 1c and 1d). We term the bottlebrush biohybrid “Quasar” (<u>qu</u>antitative <u>a</u>midation for <u>s</u>terically <u>a</u>ugmented oligodeoxyribonucleotide) for their analogous shape with the homonymous astronomical object.

**Figure 1.**
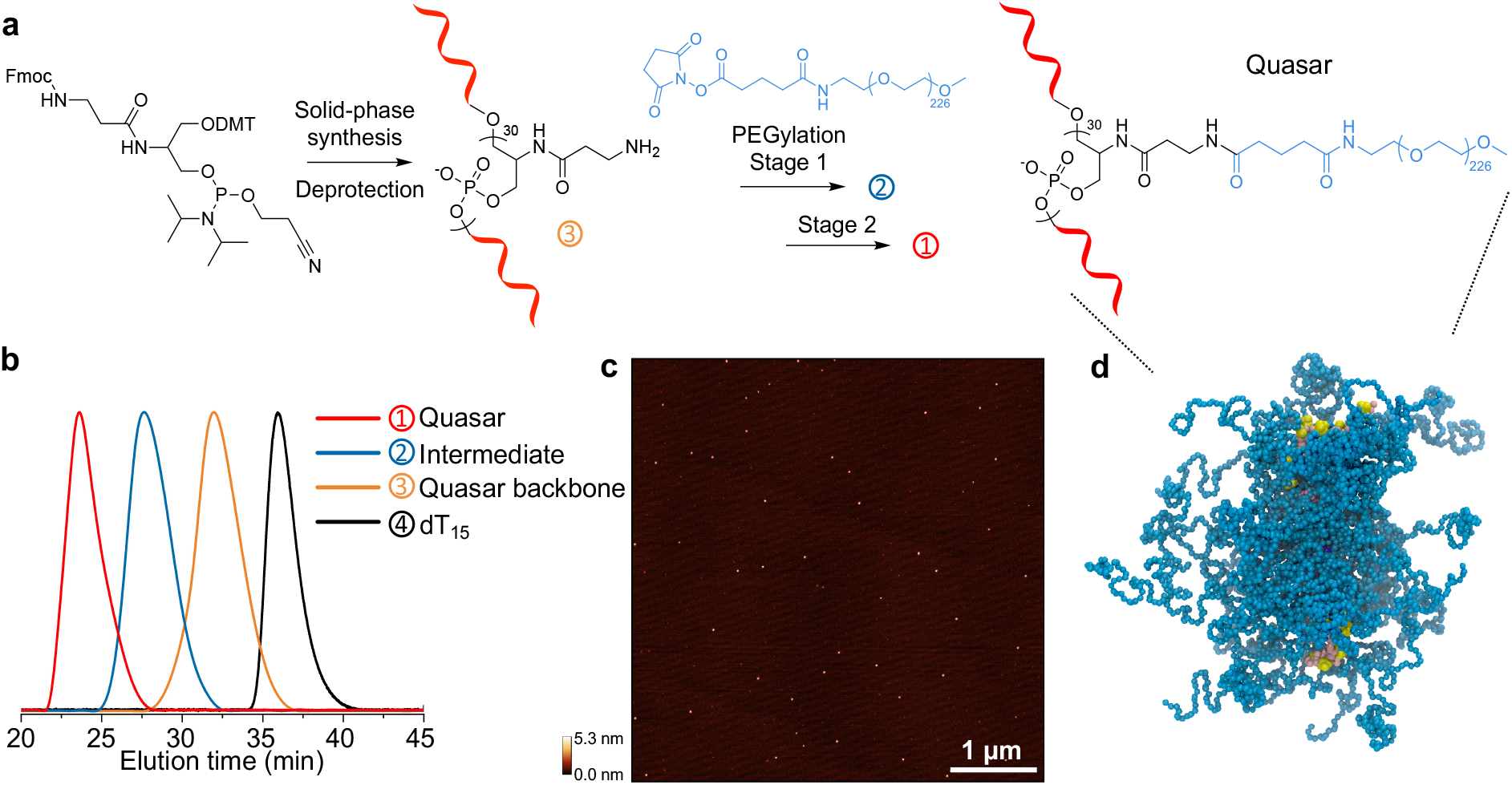
Characterization of Quasar. (a) Schematics of Quasar synthesis. (b) Aqueous GPC chromatograms of Quasar, intermediate Quasar, Quasar backbone, and free dT_15_. (c) Representative low-magnification AFM image of Quasar, showing highly homogeneous, non-aggregating spherical particles. (d) The structure of the Quasar from a coarse-grained molecule dynamics simulation using the MARTINI force field with explicit solvation (blue: PEG; yellow/pink: DNA).

The solid-phase methodology of Quasar synthesis carries significant advantages over the graft-through approach that we have adopted previously for PN bottlebrush polymers. For example, the degree of polymerization (DP) can be arbitrarily tuned by the number of synthesis cycles. Further, the synthesis provides access to additional molecular architectures, such as branched, dendritic, or block copolymers (or a combination thereof), which can be difficult to achieve with current polymer chemistry.

To test the versatility of our approach, we synthesized PSP bottlebrushes with different backbone DPs ranging from 5 to 35 (a 5’ Cy5 was used for quantification). In addition, we synthesized two architecturally distinct forms of Quasars: the Dumb-brush, where the DNA strand is situated between two bottlebrush segments, similar to a dumbbell, and the Doubler-brush, where the DNA is tethered via the 5’ to the middle unit of the brush backbone (Table S1). The Dumb-brush was synthesized with a dT_15_ bridge between two PSP_15_ segments. For the Doubler-brush, a dT_15_ segment was first synthesized normally (3’ to 5’), followed by the addition of a two-way branching unit (Doubler) at the 5’, upon which two serinol phosphoramidites were added in each coupling sequence. One benefit for the Doubler-brush is that the brush backbone can be synthesized with half of the number of synthesis cycles while achieving the same total number of repeating units. The backbone sequences were purified by RP-HPLC (Figure S1). TNBSA assay shows that the number of available amine groups associated with each backbone matches the expected value (Table S2). After the first stage of PEGylation, ∼85% of all backbone amine groups were consumed, and the yield increased to 90%-100% after the second stage (Table S2). PSP-backboned bottlebrushes of DP 5, 20, and 35 show an increase in MW and size as evidenced by aqueous GPC (Figure 2a) and dynamic light scattering (DLS) (Figure S3) measurements. Despite the different architectures and insertion positions of dT_15_, Quasar and the structural variants of similar MW exhibit similar retention times in aqueous GPC (Figure 2b) and hydrodynamic sizes (Figure S3). Narrow polydispersity indices (PDIs) in the range of 1.01-1.11 were observed for all samples. DMF GPC shows slightly higher PDIs in the range of 1.1-1.4 and larger-than-calculated MW, likely due to the polar bottlebrush backbone causing some aggregation in DMF, manifesting in high MW tailing (Figure S4). Both transmission electron microscopy (TEM, Figure S5) and AFM (Figure 2d and S6) confirm that these molecular brushes are non-aggregating and highly uniform in size, and particle size is consistent with backbone DPs. Consisting of a multitude of phosphodiesters (pKa ∼2.2), PSP bottlebrushes and Quasars exhibit negative ζ potentials ranging from -11.1 to -26.1 mV under pH neutral conditions (Nanopure™ water) (Figure 2c). Interestingly, Quasars and a PN-based counterpart of similar MW and hydrodynamic size (Figure S3) exhibit very different ζ potentials: while the PSP materials show highly negative ζ potentials similar to free oligonucleotides, the PN-based material has a near-neutral ζ potential (Figure 2c). Collectively, these results indicate that the Quasar synthesis is robust and can be used to prepare molecular brushes containing functional oligonucleotides with a high degree of structural freedom.

**Figure 2.**
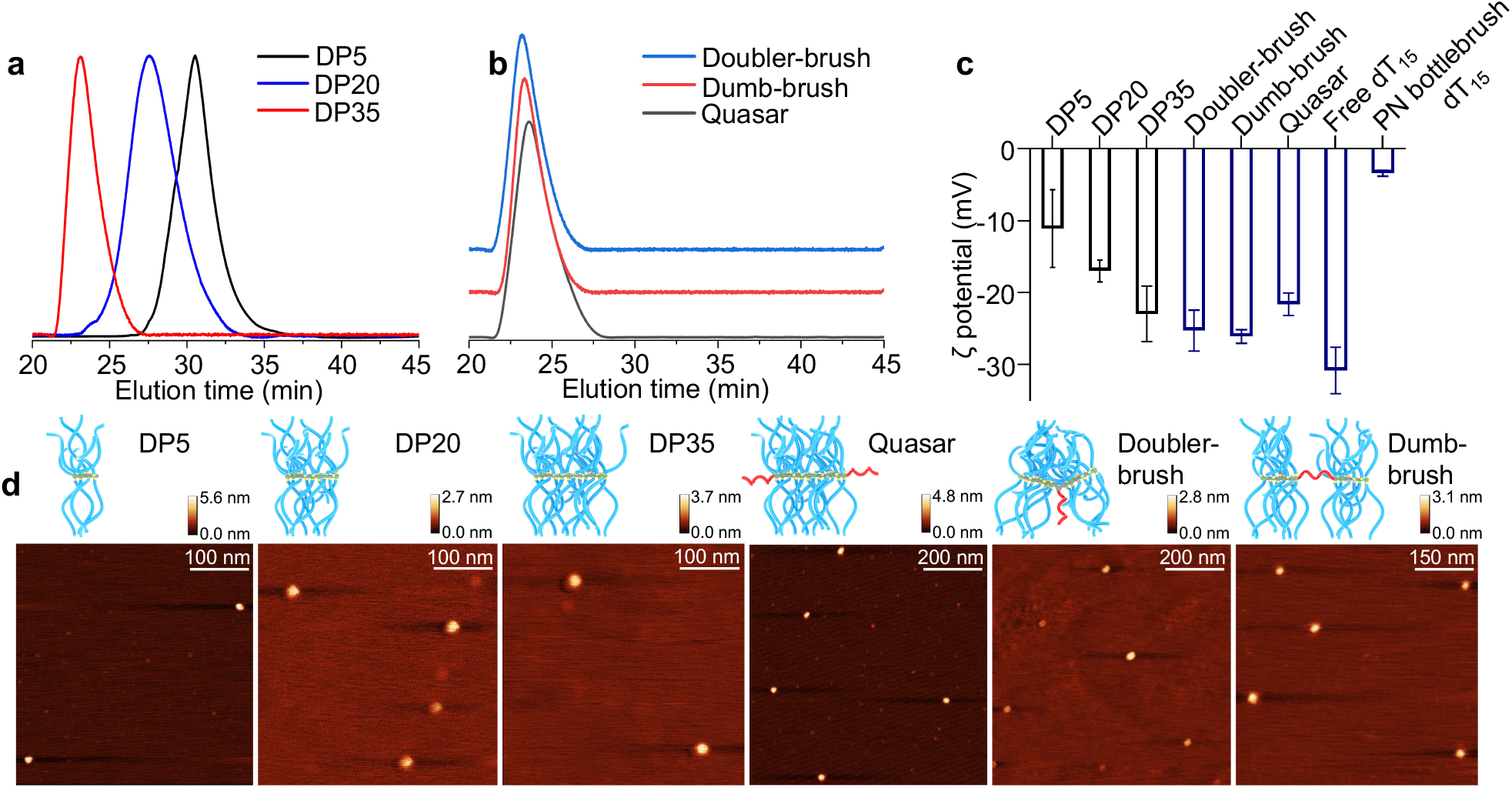
Characterizations of PSP bottlebrushes and Quasars. (a) Aqueous GPC chromatograms of the PSP bottlebrushes with DPs of 5, 20, and 35. (b) Aqueous GPC chromatograms of Quasars of various architectures. (c) Zeta potential measurements of PSP bottlebrushes, Quasars, free dT_15_ and PN bottlebrush dT_15_. (d) Representative AFM images of PSP bottlebrushes and Quasars.

The Quasars are designed to reduce unwanted oligonucleotide-protein interactions, inhibit non-specific cellular uptake, and prolong blood circulation times. To test the first characteristic, we examined the nuclease degradation kinetics of various dT_15_ Quasars (Figure S7a). The 3’ of dT_15_ was labeled with the fluorophore Cy5, and a 5’ quencher (dabcyl)-labeled complementary dA_15_ strand was hybridized to the Quasars. The hybridizing kinetics for all Quasars were indistinguishable from that of free dT_15_, reaching completion in ∼10 s. The addition of a non-complementary, quencher-labeled dummy strand did not result in a reduction in Cy5 fluorescence (Figure S7b), ruling out non-specific binding. Upon digestion by the endonuclease DNase I, the fluorophore-quencher pair is separated, leading to an increase of fluorescence. The degradation half-lives of the Quasars are roughly three times that of free double-stranded DNA, suggesting steric shielding effects (Figure S7c and Table S3). In contrast, PEGylation by conventional linear or slightly branched PEG cannot alter the degradation half-life by DNase I.^[13]^

We speculate that a mechanism for the brush-type PEG to enter the cell involves transient adsorption of the polymer onto the plasma membrane (PM), possibly mediated by PEG-cation interactions and the negative PM potential.^[14]^ With PN-based bottlebrushes, the near-neutral ζ potential promotes PM adsorption, and the hydrophobic polymer backbone can further increase adhesion strength^[15]^ and therefore polymer residence time on the PM, allowing for increased uptake compared with normal PEG or DNA. In contrast, the more negative ζ potential and the completely hydrophilic backbone of the Quasar should reduce the transient polymer-PM interactions, and therefore endocytosis would not be enhanced compared to free DNA. To test this hypothesis, we compared the cellular uptake of a Cy3-labeled Quasar and a PN-based counterpart by HUVEC (endothelial), NCI-H358 (lung), HEP3B (liver), and SKBR3 (breast) cells using flow cytometry. The results show that the Quasar consistently undergoes very limited uptake, similar to the levels exhibited by free single-stranded DNA which has been known to exhibit insignificant levels of endocytosis (Figure 3a).^[16]^ Conversely, the PN-based conjugate, which has similar MW, architecture, and PEG/DNA content as the Quasar (Figure S8), shows 6-12 times more rapid cell uptake than Quasar. These results demonstrate that a small change in the chemical composition of the backbone (no more than 5% in overall MW) can greatly alter the biological characteristics of the bottlebrush polymer.

**Figure 3.**
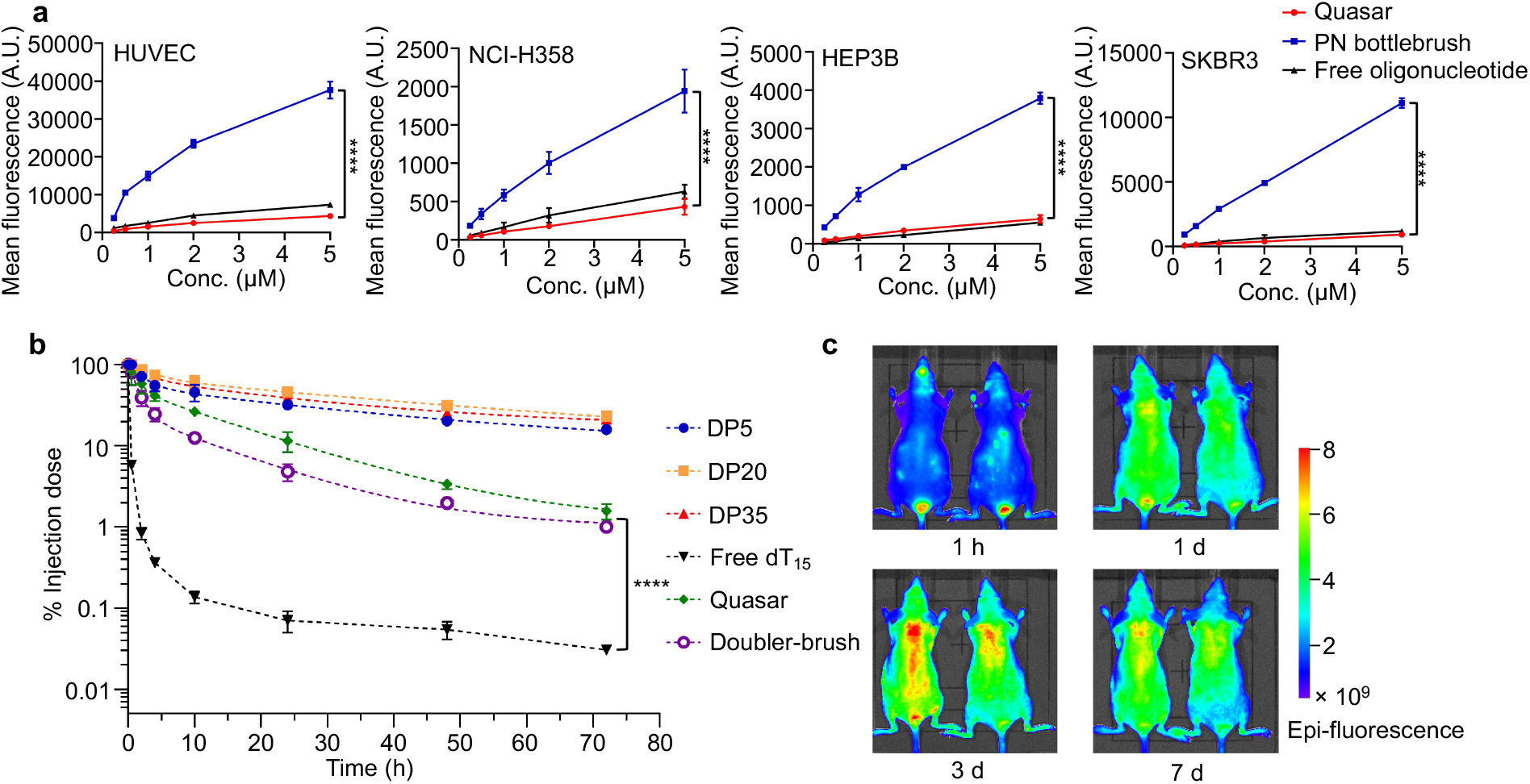
*In vitro* and *in vivo* characterization of PSP bottlebrushes and Quasars. (a) Cellular uptake of Quasar, PN-bottlebrush oligonucleotides conjugates, and free oligonucleotide by HUVEC, NCI-H358, HEP3B, and SKBR3 cells. (b) Plasma pharmacokinetics of dT_15_ Quasar, free dT_15_, Doubler-brush and PSP bottlebrushes. (c) Fluorescence imaging of SKH1-Elite mice following intravenous injection of Cy5-labeled PSP bottlebrush with a DP of 30. Statistical significance was calculated using two-way ANOVA. \*\*\*\**P*<0.0001.

Persistence in plasma is important to aptamers as rapid clearance can significantly shorten the duration of effect and increase dosage requirements. We hypothesize that Quasars can improve the plasma PK and the potency of conjugated aptamers by reducing cell uptake levels and avoiding renal clearance. To evaluate the plasma PK, 5’-Cy5-labeled PSP bottlebrushes, dT_15_ Quasar, and Doubler-brush were injected into C57BL/6 mice through the tail vein. Blood samples were collected from the submandibular vein at predetermined time points and analyzed using a two-compartment model. Remarkably, the PSP bottlebrushes exhibited very high stability and retention in plasma, with distribution half-lives in the range of 1.6 - 2.6 h and elimination half-lives between 24 and 35 h (Table S4). There was still >20% of the injected dose (for DPs = 20 and 35) remaining in circulation after 72 h (Figure 3b). For the Quasar, the plasma clearance is notably faster than the bottlebrushes alone, likely due to degradation of the dT_15_ component by plasma nucleases. The Doubler-brush was cleared slightly faster than the linear Quasar. Still, >10% of the Quasar was found within the plasma 24 h post-injection, while there was <1% free dT_15_ in circulation after only 2 h and <0.1% after 24 h. Estimating bioavailability by calculating the area under the curve (AUC_∞_), the Quasars are 15-25 times more bioavailable than free dT_15_ (Table S4).

To follow the bottlebrush component of the Quasar polymer *in vivo* for an extended period of time and investigate its biodistribution, SKH1-Elite mice, an immunocompetent hairless strain, were injected with a 5’-Cy5-labeled PSP bottlebrush with a DP of 30 through the tail vein. Live animal imaging showed that fluorescence gradually increased on the surface of the mice, reaching peak levels after 48 h. The fluorescence persisted at peak levels until approximately the 7^th^ day post injection (Figure 3c and S9), before slowly decreasing to slightly above-background levels in five weeks. On days 3, 7, and 37, animals were euthanized, and major organs were excised for imaging. All major internal organs (bar the brain) as well as muscle exhibited uptake without a strong preference for any organ. However, interestingly, pronounced accumulation in the skin (including epidermis, dermis, and the skin-draining lymph nodes) was observed. The skin was also the only organ to continue to exhibit fluorescence on day 37 (Figure S10). In immunodeficient athymic nude mice, which lack T cells, a similar observation was made (Figure S11). The skin accumulation of the Quasars may be due to their eventual uptake by skin-resident immune cells, such as dendritic cells, macrophages, and mast cells.^[17]^ Additional studies are required to elucidate the underlying mechanism for this phenomenon.

To demonstrate the binding efficacy of Quasars, we adopted a thrombin-binding aptamer, HD1, as a model system. HD1 is a 15-nucleotide DNA sequence folding into an antiparallel G-quadruplex that can specifically bind to the exosite I of human alpha thrombin (Table S1).^[18]^ Upon binding, HD1 can inhibit the coagulation and prolong coagulating times.^[19]^ Having a dissociation constant (k_d_) in the nanomolar range, HD1 has been envisioned as a short-term anticoagulant that can be used intraoperatively to reduce the risk of thromboembolism.^[20]^ Several studies have demonstrated improved *ex vivo* efficacy of HD1 through polymeric micelles or DNA origami.^[21]^ However, *in vivo* potency of HD1 has not been robustly demonstrated. We first assessed the binding affinity of HD1 and the corresponding Quasar by microscale thermophoresis (MST) using Cy5-labeled human alpha thrombin as the target. It was found that the appendage of the bottlebrush structure to HD1 has a nominal impact on its binding affinity, with free HD1 showing a k_d_ of 5.4 nM and the HD1 Quasar showing a k_d_ of 6.9 nM. In addition, a Quasar containing a scrambled sequence (Scr Quasar) exhibited no measurable binding with thrombin, ruling out the bottlebrush component being responsible for the observed binding (Figure 4a). Next, we evaluated the anticoagulation properties of HD1 and HD1 Quasar in human plasma by measuring the prothrombin time (PT) and activated partial thromboplastin time (aPTT) (Figure 4b and 4c). Interestingly, while free HD1 induced a more pronounced effect in the PT assay, the HD1 Quasar was more effective in the aPTT assay, with almost three-fold longer coagulating times compared to the vehicle. One interpretation for this difference is that the intrinsic/extrinsic pathways of the coagulation cascade are affected differently by the test agents. Again, Scr Quasar did not show anticoagulation effects. Importantly, the anticoagulation was completely reversible through the use of a locked nucleic acid (LNA)-based antidote (Table S1) that consists of a fully complementary strand to HD1, which disrupts its secondary structure.^[22]^ Such an antidote would be an invaluable tool for the perioperative management of patients receiving anticoagulants. Interestingly, although developed as a human thrombin binder, HD1 can also cause prolonged coagulation in mouse plasma, and the strengths of the effect were found to be comparable in both species. Furthermore, the diverging relative strengths of HD1 and HD1 Quasar in PT and aPTT assays were also observed in mouse plasma (Figure 5a and 5b).

**Figure 4.**
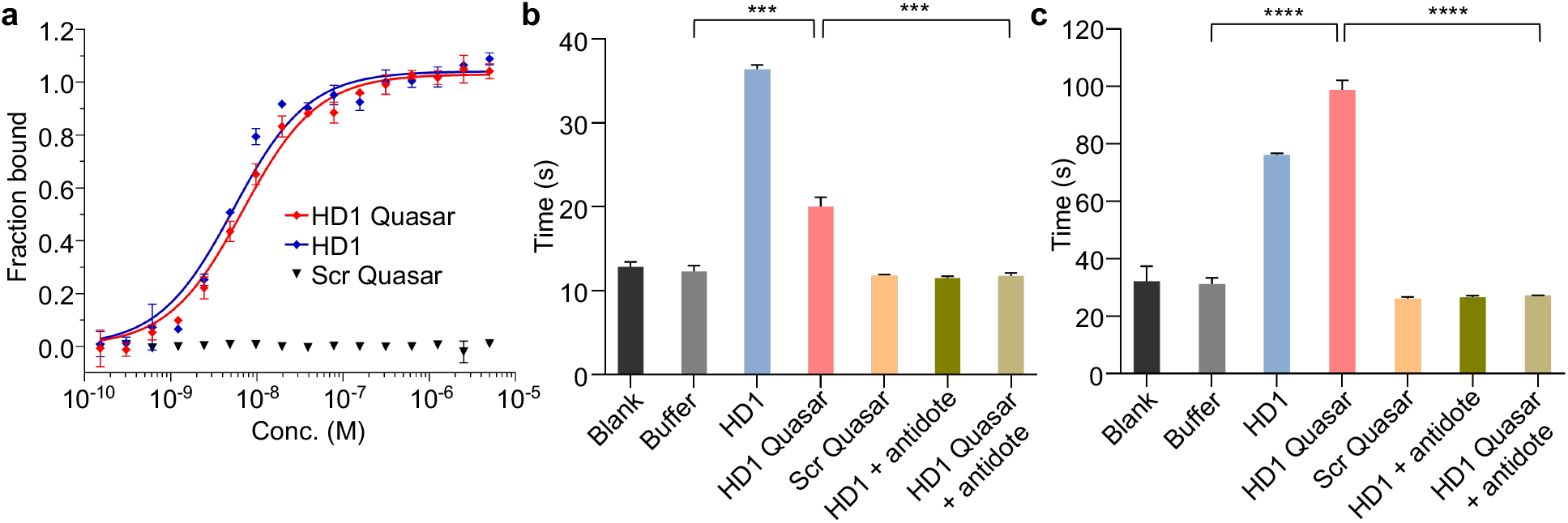
Binding affinity and anticoagulation test in human plasma. (a) Binding analysis of free HD1, HD1 Quasar, and Scramble Quasar measured by MST. (b) PT measurements of human plasma treated with samples and controls (5 μM). (c) aPTT measurements of human plasma treated with samples and controls (5 μM). Statistical significance was calculated using Student’s two-tailed t test. \*\*\**P*<0.001, \*\*\*\**P*<0.0001.

**Figure 5.**
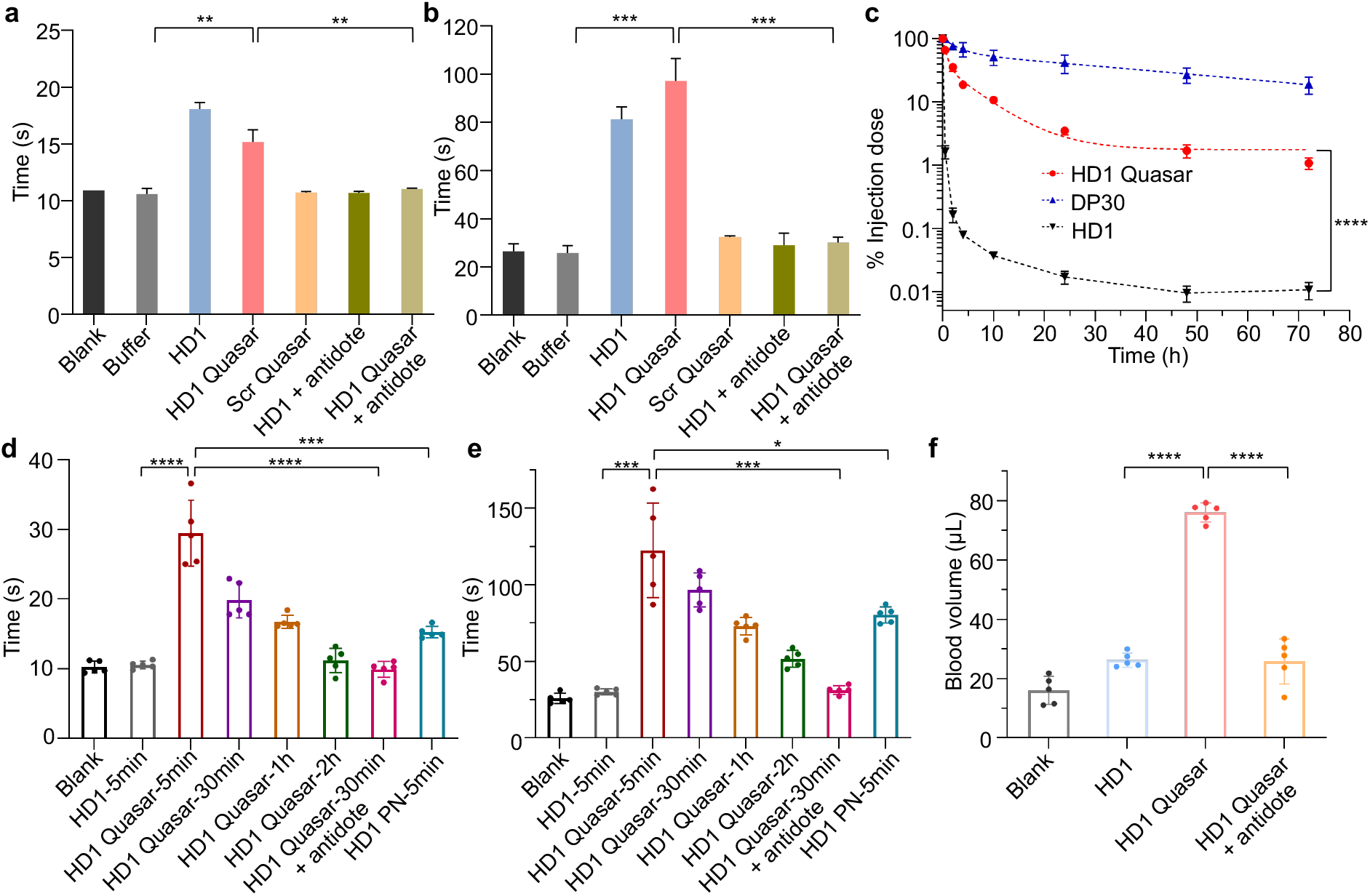
Anticoagulation by Quasar in mouse plasma and *in vivo*. Measurements of PT (a) and aPTT (b) of mouse plasma (*ex vivo*) treated with samples and controls (5 μM). Statistical significance was calculated using Student’s two-tailed *t* test. \*\**P*<0.01, \*\*\**P*<0.001. (c) Plasma pharmacokinetics of free HD1, HD1 Quasar and PSP bottlebrush (DP30). Statistical significance was calculated using two-way ANOVA. \*\*\*\**P*<0.0001. Measurements of PT (d) and aPTT (e) of mouse plasma withdrawn at predetermined time points after i.v. injection of samples and controls in C57BL/6 mice (*in vivo*). (f) Tail transection bleeding by C57BL/6 mice after treated i.v. with samples and controls. Statistical significance was calculated using Student’s two-tailed t test. \**P*<0.01, \*\**P*<0.01, \*\*\**P*<0.001, \*\*\*\**P*<0.0001.

The complex and dynamic *in vivo* environment represents the ultimate challenge for aptamers. To assess the ability of the Quasar to serve as an aptamer enhancer *in vivo*, free HD1 and the corresponding HD1 Quasar were injected intravenously in C57BL/6 mice. Plasma PK measurements of 5’-Cy5-labeled materials reveal that the HD1 Quasar persists in the blood markedly longer than free HD1, with a 16-fold difference in AUC_∞_ (Figure 5c). Interestingly, placing the Cy5 at the 5’ of the Quasar results in slightly better blood retention than placement at the 3’ (see Figure S12 for PK parameters). When analyzing anticoagulating properties, it was found that free HD1 produced no statistically significant difference compared to blank in both PT and aPTT assays using blood collected 5 min after injection, although in purified plasma HD1 does exhibit potency (*vide supra*). The rapid and complete loss of activity of free HD1 *in vivo* may be attributed to a combination of degradation and rapid renal clearance. In stark contrast, the HD1 Quasar exhibited a 3-fold increase in PT and a 5-fold increase in aPTT measurements relative to blank (Figure 5d and 5e). The effect attenuates slowly, reaching baseline levels in ∼2 h. When a dose of the LNA antidote (10 equiv. to HD1) was delivered intravenously 20 min after the administration of HD1 Quasar, the anticoagulating effect was completely neutralized in the next blood collection timepoint (30 min). Comparing the HD1 Quasar to the PN-based counterpart, it can be seen that the PN conjugate produced <50% of the effect as the HD1 Quasar at the 5 min timepoint, although both types of polymers exhibit similar plasma PK,^[11]^ which highlights the importance of the phosphodiester backbone of the Quasar. The *in vivo* anticoagulatory effect of HD1 Quasar was further quantified using a murine tail-transection bleeding model (Figure 5f). Shortly (5 min) after receiving the test agents and controls, the tails of mice were clipped and blood from the tail was collected for 15 min. Treatment with HD1 Quasar induced the largest amount of blood loss (76 μL), while free HD1- and vehicle-treated mice lost 26 μL and 16 μL blood, respectively. When mice were given Quasar and the LNA antidote sequentially, tail bleeding reverted to the baseline rate. Taken together, these results demonstrate that the Quasar structure considerably enhances the plasma PK, bioavailability, and the potency of conjugated aptamer, and the activity is controllable using an antidote.

## Conclusion

In summary, we have developed a facile route to a novel bottlebrush polymer that can be used to enhance the pharmacological properties and *in vivo* performance of aptamers. Termed Quasar, the unimolecular polymer-oligonucleotide biohybrid can be synthesized in the same step as the therapeutic sequence via the automated solid-phase methodology, followed by near-quantitative PEGylation. The synthesis is highly versatile with regard to the size and architecture of the final polymer, and does not involve heavy metal catalysts that can complicate downstream applications. Importantly, the Quasar consists only of building blocks that are recognized as safe in pharmaceutical applications, and does not involve a non-degradable, long-chain aliphatic backbone. The hydrophilic phosphodiester backbone of the Quasar resists cellular uptake, and the spatially congested PEG environment reduces non-specific binding, leading to elevated blood retention times and increased productive binding. These properties impart the Quasar superior performance in an anticoagulation mouse model compared to the free aptamer, which exhibits potency *in vitro* but no activity *in vivo*. Of note, the aptamer we adopted is not chemically modified, making it susceptible to degrading nucleases. We envision that, with advanced, chemically stabilized aptamer variants, it is possible to design Quasars with antibody-like *in vivo* potency and durability but with all the benefits associated with aptamers.

## Acknowledgements

This work was supported by the National Institute of General Medical Sciences (1R01GM121612), the National Cancer Institute (1R01CA251730), and the National Science Foundation (DMR award number 2004947). We thank Prof. Meni Wanunu for the use of AFM, Prof. Ambika Bajpayee for the help on MST and Prof. Heather Clark for the use of *in vivo* imaging system.

## Experimental Section

### Synthesis of phosphodiester-backboned molecular brushes

The PSP backbone was prepared by solid-phase synthesis using a custom phosphoramidite (Scheme S1). To graft the PEG side chains, 100 nmol of the backbone polymer and NHS-terminated 10 kDa mPEG (1:1 amine:NHS ester) were dissolved in 1 mL of PBS (pH 7.4). The mixture was shaken at 4 °C overnight and lyophilized to give a white powder, which was then redissolved in 1 mL anhydrous DMF containing 42 μL triethylamine. To this mixture, 1 equiv. of NHS-terminated PEG (dissolved in 1 mL DMF) was added in 5 aliquots (12 h between each aliquot), and the mixture was gently shaken at room temperature for 12 h after the last aliquot before being dried *in vacuo*. The PSP bottlebrush polymer was isolated by fractionation using aqueous GPC.

### Quantification of amine groups using TNBSA assay

To determine the number of amine groups before and after PEGylation, samples were dissolved in 100 μL of 0.1M sodium bicarbonate solution (pH 8.5). Then, 50 μL of 0.01% (w/v) TNBSA in 0.1 M sodium bicarbonate solution was added to each sample. For blank control, 50 μL of 0.1 M sodium bicarbonate solution without TNBSA was added. Next, the mixtures were incubated at 37 °C for 2 h. Next, 50 μL of 10% sodium dodecyl sulfate (SDS) solution and 25 μL of 1 N HCl were added to quench the reaction. Samples were then placed in a plate reader and absorbance at 335 nm was measured. The amine concentrations were calculated using a calibration curve established with a glycine standard.

### Plasma pharmacokinetics

8∼12-week-old female C57BL/6 mice were purchased from Charles River (MA, USA). Mice were randomly divided into 9 groups (n=4). Samples were intravenously administered via the tail vein at equal DNA concentrations (500 nmol/kg). Blood samples (25 μL) were collected from the submandibular vein at varying time points (30 min, 2 h, 4 h, 10 h, 24 h, 48 h and 72 h) using BD Vacutainer blood collection tubes with sodium heparin. Heparinized plasma samples were obtained by centrifugation at 3000 rpm for 20 min, and then aliquoted into a 96-well plate. The fluorescence intensities were measured on a plate reader. The amounts of agents in the blood samples were estimated using standard curves established for each sample prepared in freshly collected plasma.

### *In vivo* blood clotting

8∼10-week-old female C57BL/6 mice (Charles River, MA, USA) were randomly divided into 8 groups (n=5). Mice were administrated of samples and controls (15 nmol; DNA basis) through the tail vein. Blood samples (180 μL) collected at predetermined time points were mixed with 20 μL of 3.2% sodium citrate buffer for 5 min. Whole blood was centrifuged at 1500 g for 15 min at 4 °C. For groups that test antidote efficacy, mice were injected with an LNA antidote (150 nmol) 20 min after the initial injection of HD1 Quasar, and blood was collected 30 min after the injection of HD1 Quasar. The collected plasma samples (50 μL) were analyzed by PT and aPTT assays (see *SI*).

### Murine tail transection bleeding

8∼10-week-old female C57BL/6 mice (Charles River, MA, USA) were divided into 4 groups (n=5). Mice were injected with 200 μL of samples and controls (15 nmol; DNA basis) via the tail vein. After 5 min, the distal 1 mm of the tail was amputated. For groups testing antidote efficacy, 200 μL of an LNA antidote (150 nmol) was administered by tail vein injection immediately after the injection of the HD1 Quasar, and the clipping of the distal 1 mm of the tail was performed 10 min after the injection of the HD1 Quasar. Blood from the tail wound was collected for 15 min after tail transection, and the amount of blood loss was measured using a previously described protocol.^[23]^

---

**Scheme 1.**
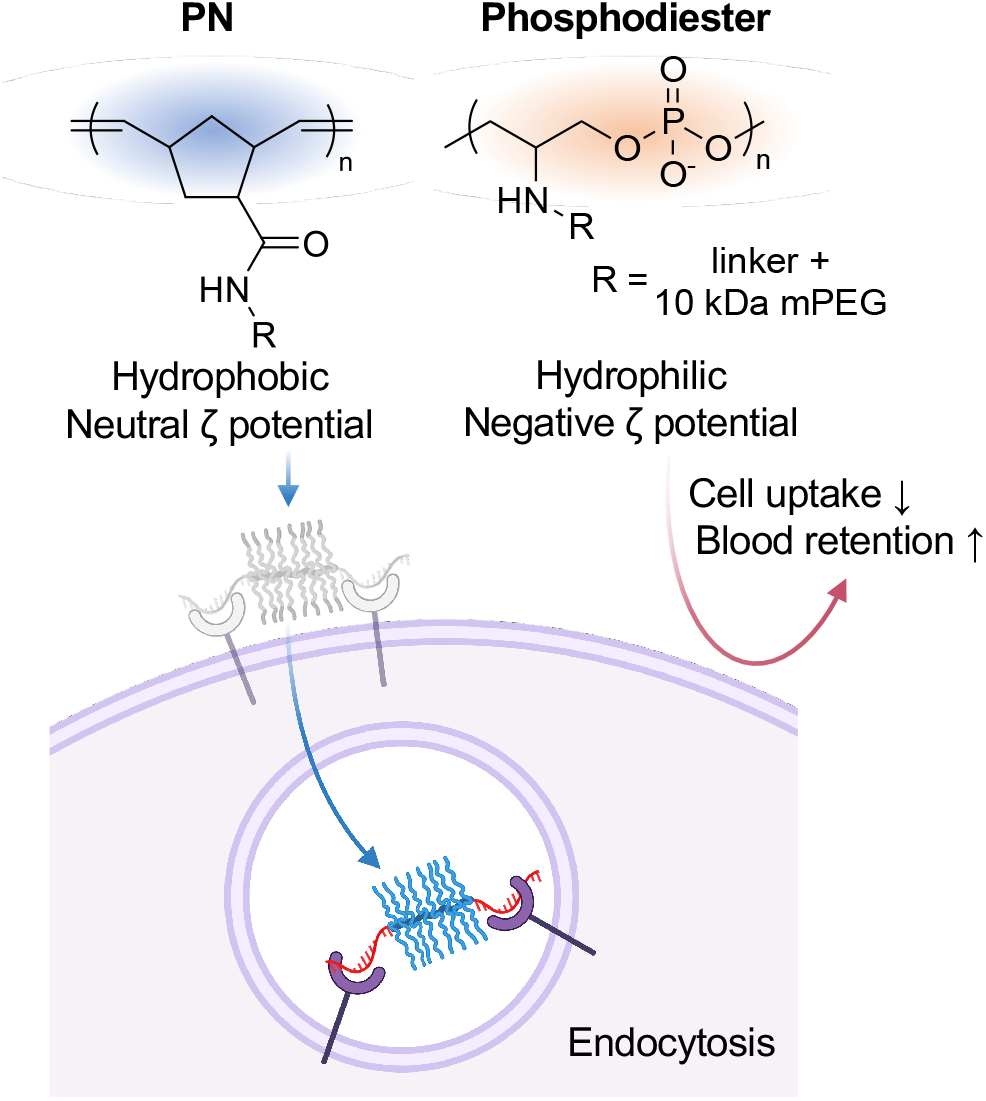
Comparison of the backbone chemistry of PN- and phosphodiester-based molecular bottlebrushes.

