## Supplementary Information for "A Long-Circulating Vector for Aptamers Based upon Polyphosphodiester-Backboned Molecular Brushes"


### Experimental Procedures

#### Materials

Methoxy polyethylene glycol (PEG) glutaramide succinimidyl ester ($M_n$=10 kDa) was purchased from Creative PEGWorks (Chapel Hill, NC, USA). Phosphoramidites and supplies for DNA synthesis were purchased from Glen Research Co. (Sterling, VA, USA). Human NCI-H358 lung cancer cell line, human SKBR3 breast cancer cell line, human Hep3B liver cancer cell line and primary human umbilical vein endothelial cell line (HUVEC) were purchased from American Type Culture Collection (Rockville, MD, USA). Human pooled normal plasma was purchased from George King Bio-Medical, Inc. (Overland Park, KS, USA). All other materials were purchased from Fisher Scientific Inc. (USA), Sigma-Aldrich Co. (USA), or VWR International LLC. (USA), and used as received unless otherwise indicated.

#### Methods

$^1$H nuclear magnetic resonance (NMR) spectra were recorded on a Varian 500 MHz NMR spectrometer (Varian Inc., CA, USA). MALDI-TOF mass spectrometry (MS) measurements were performed on a Bruker Microflex LT mass spectrometer (Bruker Daltonics Inc., MA, USA). Concentrations of samples were determined using a Nanodrop™ 2000 spectrophotometer (Thermo Scientific, USA) and BioTek Synergy Neo2 Hybrid Multi-Mode Reader (BioTek Instruments Inc., VT, USA). DLS and ζ potential measurements were performed on a Malvern Zetasizer Nano-ZSP (Malvern, UK). Samples were dissolved in Nanopure™ water at a concentration of 1 µM and filtered through a 0.2 µm PTFE filter before measurement. Fluorescence spectroscopy was carried out on a Cary Eclipse fluorescence spectrophotometer (Varian Inc., CA, USA). Reversed-phase high-performance liquid chromatography (RP-HPLC) was performed on a Waters (Waters Co., MA, USA) Breeze 2 HPLC system coupled to a Symmetry® C18 3.5 µm, 4.6×75 mm reversed-phase column and a 2998 PDA detector, using TEAA buffer (0.1 M) and HPLC-grade acetonitrile as mobile phases. Aqueous gel permeation chromatography (GPC) analysis was carried out on a Waters Breeze 2 GPC system equipped with a series of an Ultrahydrogel™ 1000, 7.8×300 mm column and three Ultrahydrogel™ 250, 7.8×300 mm columns and a 2998 PDA detector. Sodium nitrate solution (0.1 M) was used as the eluent running at a flow rate of 0.8 mL/min. The number- and weight-average molecular weights of a polymer sample were calculated based upon sodium polystyrene sulfonate (PSSNa) calibration standards with a MW range of 1,600 to 2,500 kDa (Scientific Polymer Products Inc., New York, USA), while polydispersity indices (PDIs) were determined using PAGE-purified $dT_{15}$ DNA oligomer as standard, assuming the oligomer has a PDI of 1.01. *N,N*-dimethylformamide (DMF) GPC was performed on a Tosoh EcoSEC HLC-8320 GPC system (Tokyo, Japan) equipped with a TSKGel α-M 7.8×300 mm, 13 µm column and RI/UV-Vis detectors. HPLC-grade DMF with 0.05 M lithium bromide was used as the mobile phase, and samples were analyzed at a flow rate of 0.4 mL/min. DMF-GPC calibration was based on a ReadyCal kit of polyethylene glycol standards (PSS-Polymer Standard Service-USA Inc., MA, USA). The kit covers an $M_n$ range from 232 Da to 1015 kDa. For atomic force microscopy (AFM) measurements, samples were dissolved in Nanopure$^{TM}$ water and diluted to a concentration of 1 μM. 10 μL of each sample was placed onto freshly cleaved mica (Ted Pella Inc., CA, USA) and allowed for drying. All the samples were imaged on a Dimension FastScan AFM (Bruker Corporation, USA), under the ScanAsyst in Air mode. SCANASYST-Fluid+ probes (Bruker Corporation, USA) were used for all samples. For transmission electron microscopy (TEM), samples (10 μM) were placed on parafilm as a droplet, onto which a copper-coated TEM grid was gently placed. The grids were then moved, dried, and stained using 2% uranyl acetate for 10 min. TEM images were collected on a JEOL JEM 1010 electron microscope with an accelerating voltage of 80 kV.

**Oligonucleotide and Quasar backbone synthesis**

All oligonucleotides and Quasar backbones were synthesized on a Model 391 DNA synthesizer (Applied Biosystems, Inc., Foster City, CA). The time for the coupling step of the serinol-phosphoramidite was set at 10 min (compared to 15 s for normal phosphoramidites). Oligonucleotide strands were cleaved from the CPG support and deprotected in aqueous ammonium hydroxide solution (28-30% $NH_3$ basis) at room temperature for 24 h. Quasar backbones containing Fmoc-serinol units were deprotected on-column in DMF with 20% piperidine 3× and washed with DMF 2×. The CPG was dried *in vacuo* and cleaved via the same method as normal oligonucleotide strands. All Quasar backbones and oligonucleotide strands were purified by RP-HPLC, followed by the removal of dimethoxytrityl (DMT) groups using 20% acetic acid.

**Scheme S1. Synthesis of serinol-phosphoramidite (3)**

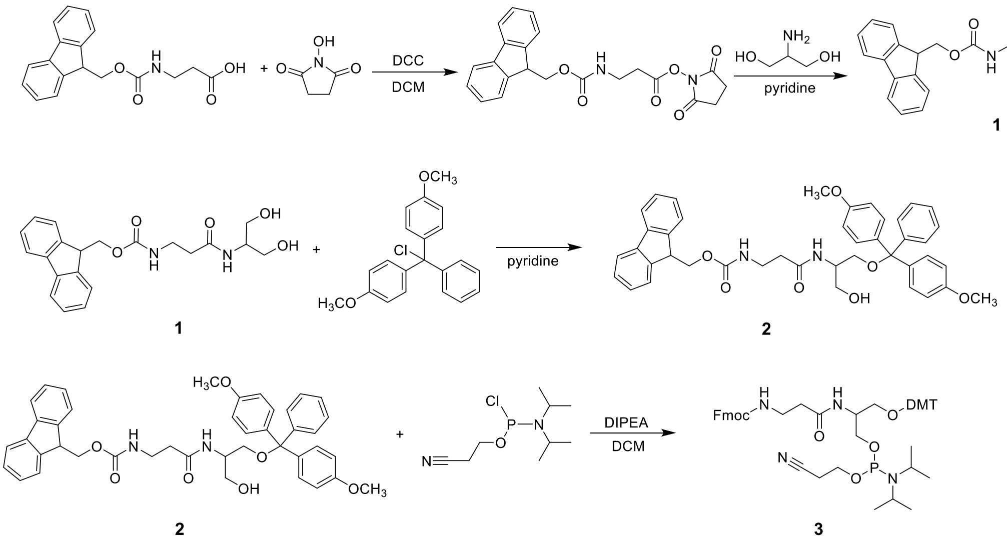


**Synthesis of Fmoc-β-Ala-serinol (1, Scheme S1)**

Fmoc-β-alanine (9.33 g, 30 mmol) and *N*-hydroxysuccinimide (3.45 g, 30 mmol) were dissolved in a mixed solvent of dichloromethane (DCM, 120 mL) and DMF (6 mL). *N,N'*-dicyclohexylcarbodiimide (DCC, 6.18 g, 30 mmol) was dissolved in DCM (12 mL) and added to the mixture. The mixture was stirred at room temperature for 1.5 h, and a white precipitate of *N,N'*-dicyclohexylurea (DCU) was observed. The reaction mixture was filtered, and the liquid was transferred to a mixture containing serinol (2.73 g, 30 mmol) and pyridine (42 mL). The reaction mixture was allowed to stir for 16 h at room temperature, before the solvent was removed by evaporation to yield a semi-solid residue (containing residue pyridine). Toluene (50 mL) was added to the residue and co-evaporated under reduced pressure 3× to remove pyridine. The resulting white solid was refluxed in DCM (200 mL) for 2 h, before being chilled to -20 °C. The product was then collected by filtration and washed with DCM (100 mL) 2× and diethyl ether (100 mL) 1×.[1] The final product was dried under high vacuum and stored at -20 °C. The isolated yield for **1** is 75±5%. $^1$H NMR (500 MHz, DMSO-$d_6$) δ 7.89 (d, J = 7.6 Hz, 2H), 7.68 (d, J = 7.5 Hz, 2H), 7.56 (d, J = 8.2 Hz, 1H), 7.42 (t, J = 7.5 Hz, 2H), 7.33 (t, J = 7.4 Hz, 2H), 7.27 (t, J = 5.9 Hz, 1H), 4.58 (t, J = 5.7 Hz, 2H), 4.27 (d, J = 6.7 Hz, 2H), 4.20 (t, J = 7.1 Hz, 1H), 3.70 (q, J = 6.3 Hz, 1H), 3.38 (t, J = 5.4 Hz, 4H), 3.18 (q, J = 6.9 Hz, 2H), 2.27 (t, J = 7.3 Hz, 2H); Maldi-TOF: m/z calculated for $C_{21}H_{24}N_2O_5$ [M-H]$^-$ 383.17, found 383.70.

**Synthesis of Fmoc-β-ala-serinol-DMT (2)**

Fmoc-β-Ala-serinol (**1**, 5.88 g, 15.3 mmol) was placed in a flask filled with $N_2$, and was dissolved in pyridine (16 mL). The flask was chilled in an ice bath. 4,4'-DMT chloride (5.37 g, 15.9 mmol) was dissolved in pyridine (32 mL) and added dropwise to the mixture containing **1**. The mixture was allowed to stir under $N_2$ for 1 h in an ice bath and then overnight at room temperature. Methanol (1 mL) was added to the mixture and stirred for 15 min to quench the reaction. Removal of the solvent under reduced pressure yields an oily residue, which was then co-evaporated with toluene (40 mL) 3×. The mixture was dissolved in DCM (50 mL) and washed with 5% sodium bicarbonate solution (50 mL) and then with brine (50 mL), before being dried over anhydrous sodium sulfate. The crude product was concentrated and purified by silica gel column purification (ethyl acetate/methanol/triethylamine 95:5:1 v:v:v). The isolated yield of **2** is 43±5%. $^1$H NMR (500 MHz, Chloroform-*d*) δ 7.75 (d, J = 7.5 Hz, 2H), 7.57 (dd, J = 7.4, 2.5 Hz, 2H), 7.39 (d, J = 7.6 Hz, 4H), 7.33 – 7.24 (m, 9H), 6.83 (d, J = 8.4 Hz, 4H), 5.99 (d, J = 7.8 Hz, 1H), 5.53 (d, J = 6.5 Hz, 1H), 4.34 (dd, J = 7.3, 3.7 Hz,

2H), 4.23 – 4.14 (m, 1H), 3.81 (dd, J = 11.4, 4.8 Hz, 1H), 3.76 (s, 7H), 3.69 (dd, J = 11.3, 4.3 Hz, 1H), 3.48 (p, J = 7.3 Hz, 2H), 3.39 – 3.25 (m, 2H), 2.39 (q, J = 6.0 Hz, 2H); Maldi-TOF: m/z calculated for $C_{42}H_{42}N_2O_7$ $[M+Na]^+$ 709.29, found 709.17.

**Synthesis of serinol-phosphoramidite (3)**

Fmoc-β-Ala-serinol-DMT (**2**, 2.76 g, 4 mmol) was placed in a flask charged with $N_2$ and dissolved in dry DCM (12 mL) containing *N,N*-diisopropylethylamine (DIPEA, 3.5 mL). The flask was chilled in an ice bath. 2-Cyanothyl-*N,N*-diisopropylchlorophosphoramidite (1.9 g, 8 mmol) was dissolved in DCM (4 mL) and added dropwise to the mixture containing **2**. The reaction mixture was allowed to stir vigorously for 20 min before being warmed to room temperature and then stirred for another 40 min. An excess of ethyl acetate (EA) was added to the reaction mixture. The mixture was washed with saturated sodium bicarbonate solution (10 mL).[2] After drying over anhydrous sodium sulfate, the mixture was filtered, and the filtrate was concentrated for silica gel column purification (hexane/ethyl acetate/triethylamine 67:33:1 v:v:v). The isolated yield of **3** is 66±5%. $^1$H NMR (500 MHz, Chloroform-*d*) δ 7.77 (d, *J* = 7.5 Hz, 2H), 7.59 (dd, *J* = 7.6, 4.1 Hz, 2H), 7.46 – 7.37 (m, 4H), 7.32 (d, *J* = 7.8 Hz, 8H), 7.23 (d, *J* = 7.4 Hz, 1H), 6.84 (d, *J* = 8.2 Hz, 4H), 5.84 (d, *J* = 8.6 Hz, 1H), 5.57 (q, *J* = 6.7 Hz, 1H), 4.34 (d, *J* = 7.4 Hz, 2H), 4.16 (dt, *J* = 29.2, 7.3 Hz, 1H), 3.80 (d, *J* = 11.5 Hz, 9H), 3.71 – 3.62 (m, 2H), 3.61 – 3.51 (m, 2H), 3.49 (s, 2H), 3.42 – 3.28 (m, 1H), 3.16 (dt, *J* = 9.3, 6.2 Hz, 1H), 2.60 – 2.47 (m, 2H), 2.38 (dd, *J* = 13.3, 6.7 Hz, 2H), 1.20 – 1.11 (m, 12H).

**Scheme S2. Synthesis of polynorbornene (PN)-backboned bottlebrush oligonucleotides conjugates**

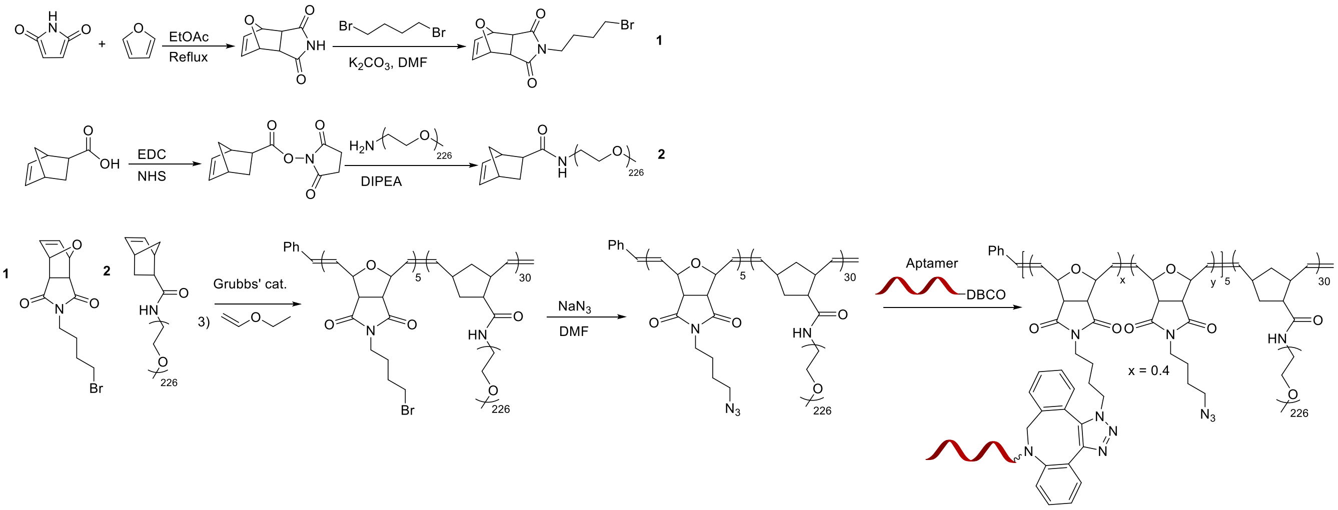


**Synthesis of PN-backboned bottlebrush oligonucleotides conjugates**

Norbornenyl bromide and norbornenyl PEG were synthesized as previously described.[3] Modified 2nd generation Grubbs catalyst was prepared based on a published method shortly prior to use.[4] Next, norbornenyl bromide (5 equiv.) was dissolved in deoxygenated DCM under $N_2$ and cooled to -20 °C in an ice-salt bath. The modified Grubbs' catalyst (1 equiv.) in deoxygenated DCM was added to the solution via a gastight syringe, and the solution was stirred vigorously for 30 min. After thin-layer chromatography (TLC) confirmed the complete consumption of the monomer, norbornenyl PEG (50 equiv.) in deoxygenated DCM was added to the reaction, and the mixture was stirred for 6 h. Several drops of ethyl vinyl ether were added to quench the reaction and the solution was stirred for an additional 2 h. After concentration under vacuum, the residue was precipitated into cold diethyl ether 3×. The precipitant was dried under vacuum to afford a white powder. Subsequently, the PN-backboned brush polymer was reacted with an excess of sodium azide in anhydrous DMF overnight at room temperature. The materials were transferred to a dialysis tubing (MWCO, 10 kDa), dialyzed against Nanopure™ water for 24 h, and lyophilized to afford a white powder. The azide-functionalized PN bottlebrush polymer (50 nmol) was dissolved in 1 mL sodium chloride solution (3 M) and reacted with dibenzocyclooctyne (DBCO)-modified oligonucleotides (100 nmol) at 50 °C overnight. The conjugate was purified by aqueous GPC, desalted, and lyophilized. The purified PN bottlebrush oligonucleotide conjugates were stored at -20 °C before use.

**Hybridization kinetics**

Samples were dissolved in PBS (pH 7.4) at a final DNA concentration of 100 nM. A total of 1 mL solution for each sample was transferred to a quartz cuvette. Dabcyl-labeled complementary strand or dummy strand (2 equiv.) in 2 μL PBS solution were added into the cuvette and rapidly mixed with a pipette. The fluorescence of the solution (ex = 640 nm, em = 670 nm) was continuously monitored every 3 seconds for 30 min. The endpoint was determined by adding a large excess (10 equiv.) of the complementary dabcyl strand to the mixture. The kinetics plots were normalized to the endpoint determined for each sample, and all measurements were repeated 3×.

**Coarse-grained molecular dynamics simulations.** An all-atom structure of the polymer biohybrid with the poly(serinol phosphodiester) backbone was mapped to coarse-grained (CG) beads according to functional groups that best match the bead types in the MARTINI force field[5] (2-5 atoms per bead). The CG parameters for the polymer backbone and linkers were extracted from a molecular dynamics (MD) trajectory of an atomistic simulation of a three-repeating unit model molecule based on the OPLS-AA force field.[6] The coarse-grained structure was solvated in a CG water box. Sodium ions were added to ensure the system is neutral in charge. The solvated system underwent energy minimization, followed by 50 ns of equilibration and 1 μs of production MD simulation (step size: 4 fs; NPT ensemble) using GROMACS 2021.3[7] with the velocity rescale thermostat[8] and the Parrinello-Rahman barostat[9] under 300 K and 1 bar.

**Nuclease degradation kinetics**

Samples were each mixed with their complementary dabcyl-labeled DNA (2 equiv.) in PBS. The solutions were heated to 95 °C for 5 min and cooled down to room temperature, then shaken overnight. Next, 100 µL of each sample was withdrawn and diluted to 1 mL (100 nM) with assay buffer (10 mM Tris-HCl, 2.5 mM $MgCl_2$, and 0.5 mM $CaCl_2$, pH 7.5). The mixture was transferred to a quartz cuvette which was mounted on a fluorimeter. DNase I was added and rapidly mixed to give a final concentration of 0.6 unit/mL. The fluorescence of the samples (ex = 640 nm, em = 670 nm) was measured immediately and every 3 seconds for 2 h. The endpoint was determined by adding a large excess of DNase I (5 units/mL) to the solution followed by incubation for 2 h.

**Cell culture**

NCI-H358 and SKBR3 cells were cultured in RPMI 1640 media supplemented with 10% fetal bovine serum (FBS) and 1% antibiotics. Hep3B cells were cultured in DMEM media supplemented with 10% FBS and 1% antibiotics. HUVEC cells were cultured in endothelial cell basal medium 2 (PromoCell, Germany) and supplemented with SupplementPack endothelial cell GM2 (PromoCell, Germany). Cell culture flasks and well plates for HUVEC were coated with 0.1% gelatin overnight before use. All cells were cultured at 37 °C in a humidified atmosphere containing 5% $CO_2$.

**Cellular uptake**

Cellular uptake of samples and controls was evaluated using flow cytometry. Cells were seeded in 24-well plates at a density of $2.0\times10^5$ cells per well in 1 mL full growth media and cultured for 24 h at 37 °C with 5% $CO_2$. After washing by PBS 1×, Cy3-labeled samples and controls (250 nM – 5 µM equiv. of DNA) dissolved in serum-free culture media (400 µL) was added, and cells were further incubated at 37 °C for 4~6 h. Next, cells were washed with PBS 2× and treated with trypsin (60 µL per well). Thereafter, 1 mL of PBS was added to each culture well to suspend the cells. Cells were then analyzed on an Attune™ NxT flow cytometer (Invitrogen, MA). Data for $1.0\times10^4$ gated events were collected.

**Biodistribution**

Animal protocols were approved by the Institutional Animal Care and Use Committee of Northeastern University. Animal experiments and operations were conducted in accordance with the approved guidelines. 8~10-week-old female athymic mice and SKH1-Elite mice were purchased from Charles River (MA, USA). Cy5-labeled samples (10 nmol in 200 µL PBS) were injected into mice through the tail vein. Fluorescent images were collected at 1, 4, 8, 24 h and daily thereafter using an IVIS Lumina II imaging system (Caliper Life Sciences Inc., MA, USA). At predetermined time points, mice were euthanized using $CO_2$. Major organs (heart, liver, spleen, lung, kidney, brain, skin) were dissected and rinsed with PBS for biodistribution analysis.

**Microscale thermophoresis**

Microscale thermophoresis (MST) binding measurements were carried out with 10 nM Cy5-labeled human alpha thrombin as target. On average there were 2.5 Cy5 dyes per thrombin protein. Samples and controls were dissolved in binding buffer (20 mM HEPES, pH 7.4, 150 mM NaCl, 2 mM $CaCl_2$, and 0.05% TWEEN) at 10 µM as stock solutions. Then, 10 µL of each sample was serial-diluted (halving concentration each time) for a total of 16 dilutions, and each dilution was mixed with 10 µL of 20 nM Cy5-labeled human alpha thrombin solution. The mixtures were transferred to Monolith NT.115 standard capillaries and analyzed on a Monolith NT.115 instrument (NanoTemper Technologies, Munich, Germany) at medium MST power and 20% excitation power. Data were analyzed using MO. Affinity Analysis software (version 2.3, NanoTemper Technologies) and MST-on time was set at 1.5 s.

**Plasma clotting assays using human and mouse plasma**

Prothrombin time (PT) and activated partial thromboplastin time (aPTT) assays were performed on a model BFT-2 coagulometer (Siemens, USA). Whole blood from C57BL/6 mice was collected in a tube containing sodium citrate solution (3.2% w/v) to a ratio of ~9:1 (blood:citrate solution). Plasma was collected after centrifuging at 1500 g for 15 min. For PT, normal human plasma or mouse plasma (50 µL) were mixed with 5 µL of samples or controls to give a final DNA concentration of 5 µM. To test antidote efficacy, 5 µL of antidote (10 equiv. of complementary DNA) was added to the plasma. The samples were incubated at 37 °C for 5 min, to which 100 µL of thromboplastin-D (ThermoFisher, MA, USA) was added to initiate the coagulation. The time until clot formation was automatically recorded by the coagulometer. For aPTT, normal human plasma or mouse plasma (50 µL) was mixed with 5 µL of samples and controls to give a final DNA concentration of 5 µM. To test antidote efficacy, 5 µL of antidote (10 equiv. of complementary DNA) was added to the plasma. The samples were then incubated with 50 µL of aPTT-XL (ThermoFisher, MA, USA) at 37 °C for 5 min before 50 µL of $CaCl_2$ (0.025 M) was added to initiate the coagulation. The time until clot formation was automatically recorded by the coagulometer.

**Table S1.** All Quasar structures and sequences used in this study.

| Sample ID | Sequence |
|---|---|
| **DP5** | 5'-Cy5-XXX XXT-3' |
| **DP20** | 5'-Cy5-XXX XXX XXX XXX XXX XXX XXT-3' |
| **DP30** | 5'-Cy5-XXX XXX XXX XXX XXX XXX XXX XXX XXX XXX T-3' |
| **DP35** | 5'-Cy5-XXX XXX XXX XXX XXX XXX XXX XXX XXX XXX XXX XXT-3' |
| **$dT_{15}$ Quasar** | 5'-Cy5-TTT TTT TTT TTT TTT XXX XXX XXX XXX XXX XXX XXX XXX XXX XXX TTT TTT TTT TTT TTT-3' |
| **Doubler-brush** | (5'-XXX XXX XXX XXX XXX$)_2$ D TTT TTT TTT TTT TTT-Cy5-3' |
| **Dumb-brush** | 5'-Cy5-XXX XXX XXX XXX XXX TTT TTT TTT TTT TTT XXX XXX XXX XXX XXX T-3' |
| **$dT_{15}$** | 5'-TTT TTT TTT TTT TTT-Cy5-3' |
| **Quencher** | 5'-Dabcyl-AAA AAA AAA AAA AAA-3' |
| **Dummy quencher** | 5'-Dabcyl-TTT TTT TTT TTT TTT-3' |
| **HD1** | 5'-GGT TGG TGT GGT TGG-3' |
| **HD1-DBCO** | 5'-DBCO-TTT TTG GTT GGT GTG GTT GG-3' |
| **HD1 Quasar** | 5'-GGT TGG TGT GGT TGG TTT TTX XXX XXX XXX XXX XXX XXX XXX XXX XXX XXT TTT TGG TTG GTG TGG TTG G -3' |
| **Scramble HD1 Quasar** | 5'-GGT GGT GGT TGT GGT TTT TTX XXX XXX XXX XXX XXX XXX XXX XXX XXX XXT TTT TGG TGG TGG TTG TGG -3' |
| **Antidote** | 5'-CCA ACC ACA CCA ACC-3' |
| **Cy3-Quasar** | 5'-Cy3-GCT ATT AGG AGT CTT TXX XXX XXX XXX XXX XXX XXX XXX XXX XXX XT-3' |
| **Cy3-oligonucleotide-DBCO** | 5'-DBCO-GCT ATT AGG AGT CTT T-Cy3-3' |
| **Cy3-oligonucleotide** | 5'-GCT ATT AGG AGT CTT T-Cy3-3' |
| **5'-Cy5-HD1** | 5'-Cy5-GGT TGG TGT GGT TGG-3' |
| **5'-Cy5-HD1 Quasar** | 5'-Cy5-GGT TGG TGT GGT TGG TTT TTX XXX XXX XXX XXX XXX XXX XXX XXX XXX XXT-3' |
| **3'-Cy5-HD1** | 5'-GGT TGG TGT GGT TGG-Cy5-3' |
| **3'-Cy5-HD1 Quasar** | 5'-XXX XXX XXX XXX XXX XXX XXX XXX XXX XXX TTT TTG GTT GGT GTG GTT GG-Cy5-3' |

X: amine-serinol phosphoramidite (to be PEGylated); D: symmetric doubler phosphoramidite; DBCO: dibenzo-cyclooctyne phosphoramidite; underlined letters indicate LNA-modified bases.

**Table S2.** The number of free amine groups on PSP backbones and the extent of PEG derivatization after each stage as determined by TNBSA assay.

| Sample ID | Amine # before PEGylation | Remaining amine/polymer after $1^{st}$ Stage | Coupling efficacy | Remaining amine/polymer after $2^{nd}$ Stage | Coupling efficacy |
|---|---|---|---|---|---|
| **DP5** | 5.3(±1.2) | 0.5 (±0.2) | 90.9% (±4.8) | Not detected | 100% |
| **DP20** | 19.5(±1.0) | 2.8 (±0.7) | 86.0% (±3.6) | 1.9 (±0.2) | 90.6% (±0.9) |
| **DP35** | 32.2(±3.0) | 5.3 (±0.6) | 85.0% (±1.7) | 2.0 (±0.4) | 94.3% (±1.2) |
| **Quasar** | 29.3(±1.8) | 4.6 (±2.0) | 84.7% (±6.6) | 1.6 (±0.2) | 94.7% (±0.7) |
| **Douber-brush** | 26.8(±1.2) | 3.6 (±0.1) | 88.0% (±0.5) | 1.5 (±0.6) | 94.9% (±2.0) |
| **Dumb-brush** | 30.0(±4.0) | 5.4 (±2.0) | 84.6% (±3.2) | 1.5 (±0.1) | 95.2% (±0.4) |

**Table S3.** Enzymatic half-lives of free $dT_{15}$ and Quasars.

| Sample ID | Free $dT_{15}$ | Quasar | Doubler-brush |
|---|---|---|---|
| $t_{1/2}$ (min) | 13.93 (±0.53) | 32.18 (±6.35) | 38.15 (±1.79) |

**Table S4.** Plasma pharmacokinetic parameters of free $dT_{15}$, PSP bottlebrushes, and $dT_{15}$ Quasars in C57BL/6 mice.

| Sample ID | $t_{1/2\alpha}$ (h) | $t_{1/2\beta}$ (h) | $AUC_{\infty}$ (nmol/ml·h) |
|---|---|---|---|
| Free $dT_{15}$ | 0.26 | 0.51 | 2.68 |
| Quasar | 1.06 | 10.80 | 67.79 |
| Doubler-brush | 1.02 | 9.47 | 38.96 |
| DP5 | 1.61 | 24.77 | 190.60 |
| DP20 | 2.63 | 35.34 | 310.56 |
| DP35 | 2.06 | 24.09 | 314.04 |

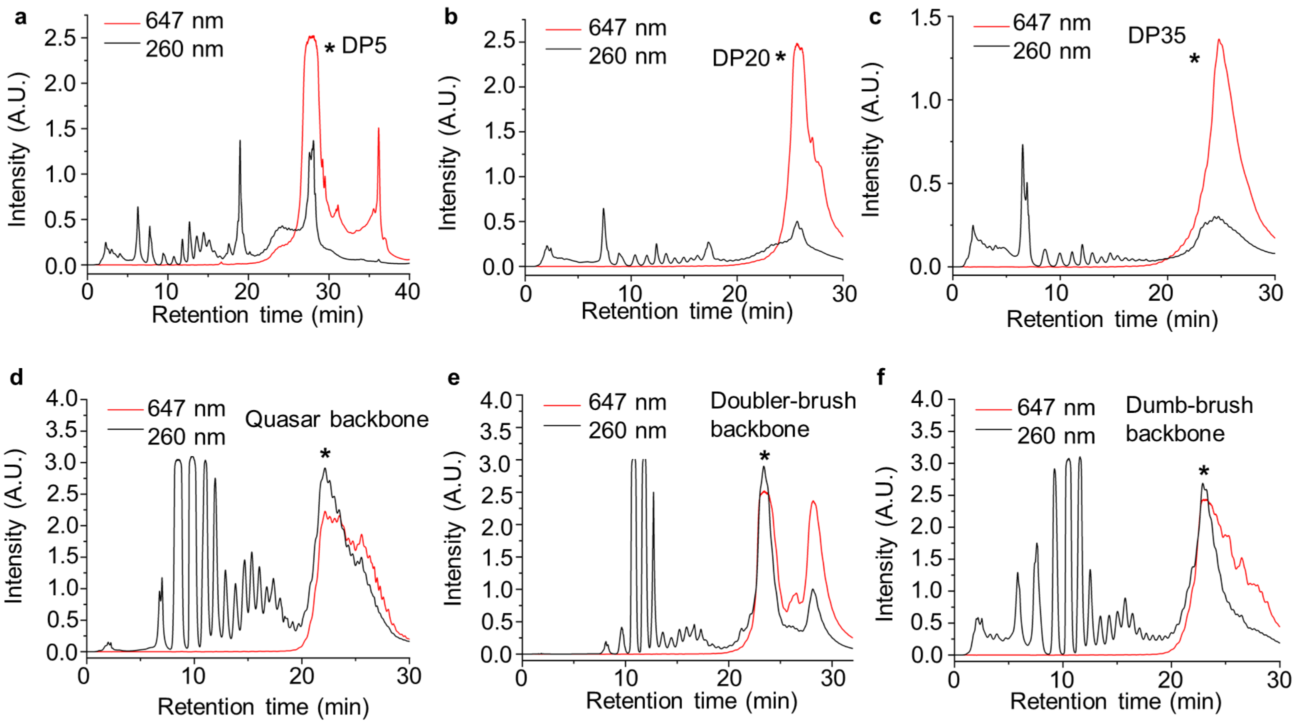


**Figure S1.** RP-HPLC chromatograms of as-synthesized PSP backbones. The peaks marked with asterisk were fractionated for further reaction.

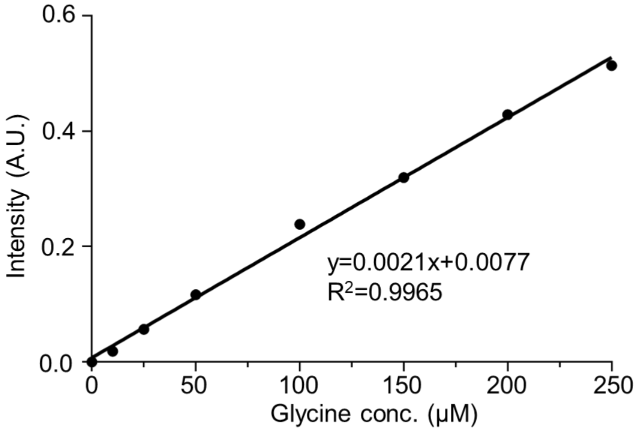


**Figure S2.** Calibration curve established using glycine as a standard for the TNBSA assay of free primary amines.

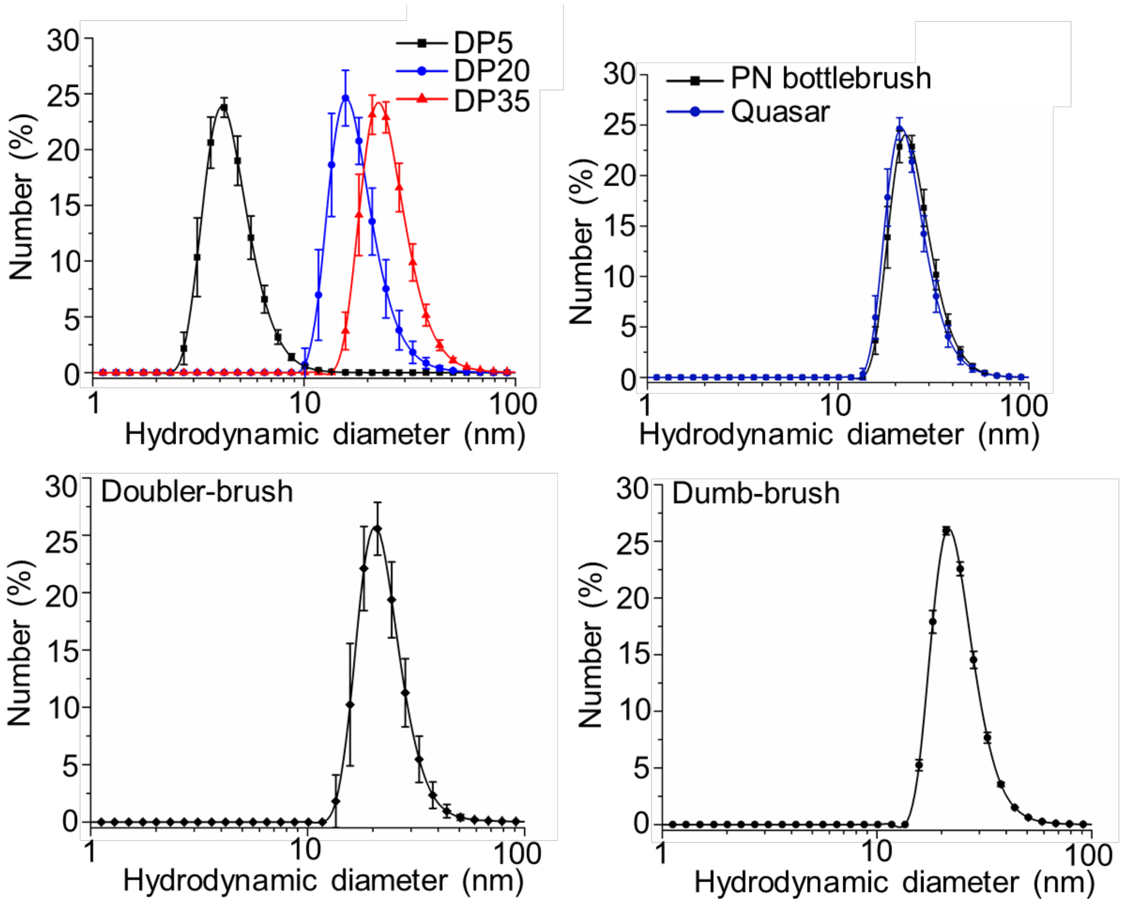


| Sample ID | DP 5 | DP 20 | DP 35 | Quasar | Doubler-brush | Dumb-brush | PN bottlebrush |
|---|---|---|---|---|---|---|---|
| **Hydrodynamic diameter (nm)** | 4.6±0.2 | 18.1±1.4 | 25.7±1.0 | 24.6±1.0 | 22.8±1.5 | 24.3±0.1 | 25.7±0.9 |

**Figure S3.** DLS measurements of PSP bottlebrushes (DP5, DP20 and DP35), PN bottlebrush, Quasar, Doubler-brush and Dumb-brush in Nanopure™ water.

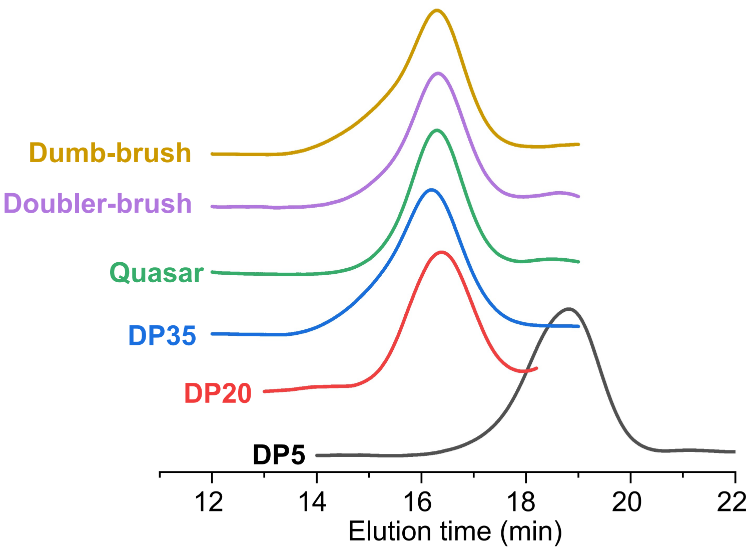


| Sample ID | $M_n$ (kDa) | $M_w$ (kDa) | PDI |
|---|---|---|---|
| **DP5** | 48.9 | 68.1 | 1.39 |
| **DP20** | 347.9 | 414.9 | 1.19 |
| **DP35** | 421.8 | 582.9 | 1.38 |
| **Quasar** | 374.8 | 448.0 | 1.19 |
| **Doubler-brush** | 374.6 | 442.5 | 1.18 |
| **Dumb-brush** | 410.8 | 547.9 | 1.33 |

**Figure S4.** DMF-GPC chromatogram (top) and average MW (bottom) of PSP bottlebrushes and Quasars.

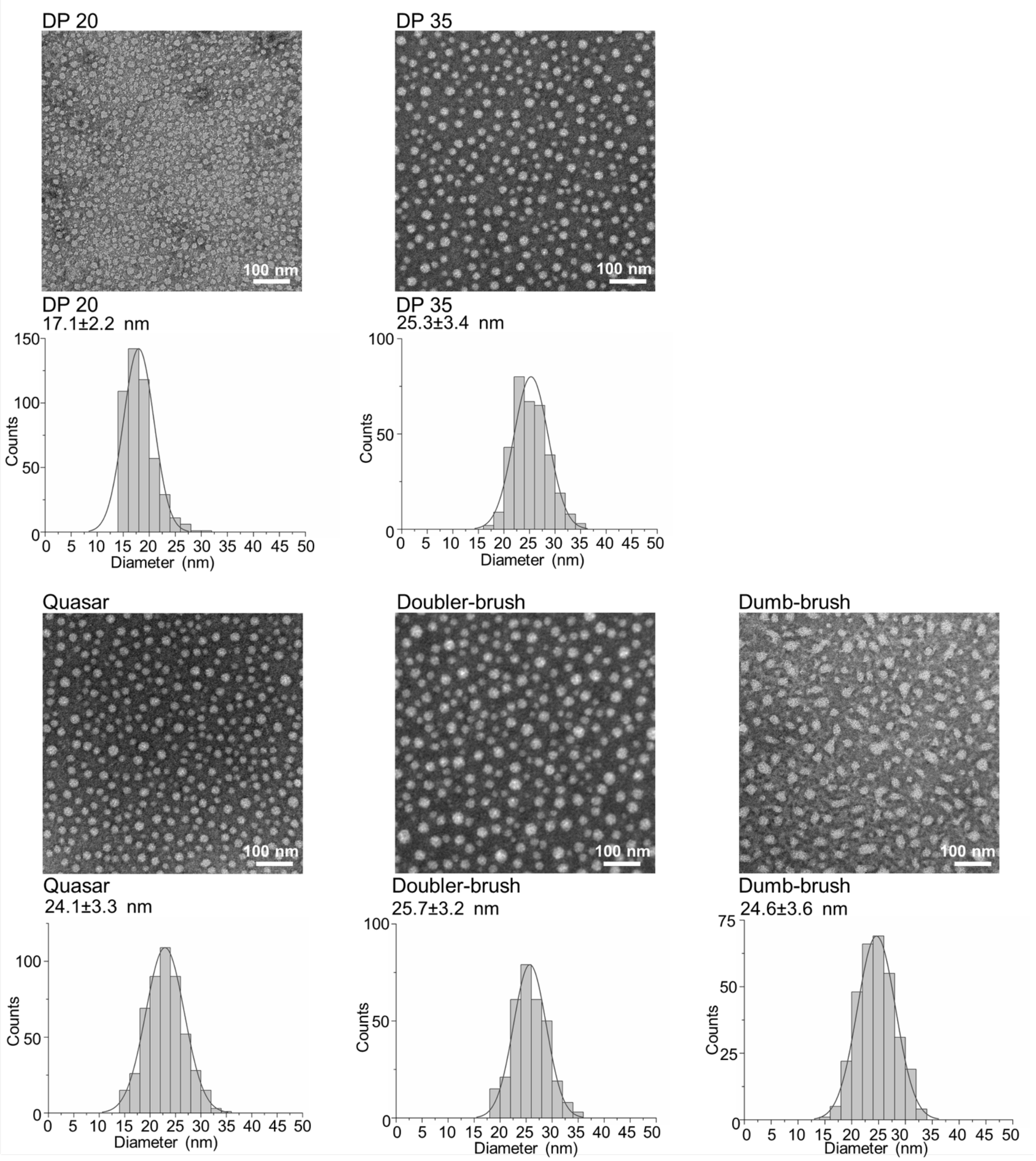


**Figure S5.** TEM images (negative stain using 2% uranyl acetate) and size measurements of PSP bottlebrushes (DP20 and DP35), Quasar, Doubler-brush and Dumb-brush. A minimum of 300 particles per sample were measured.

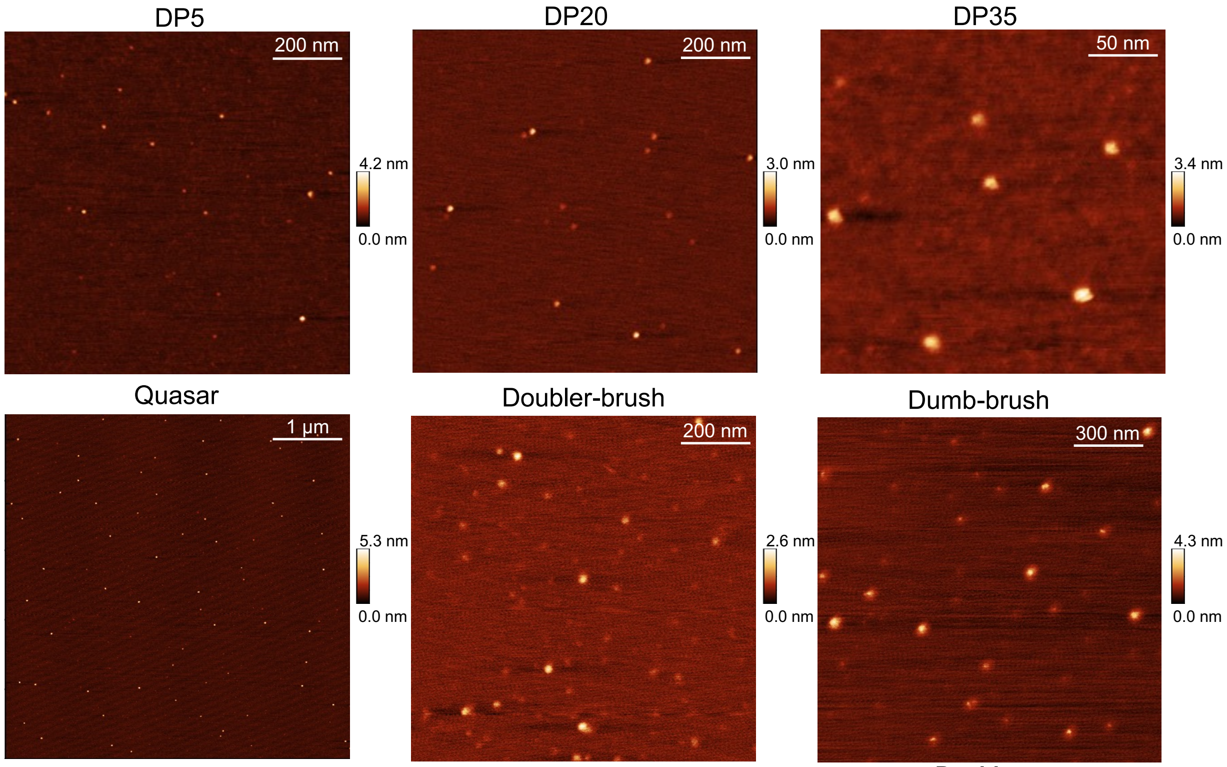


| **Sample ID** | **DP5** | **DP20** | **DP35** | **Quasar** | **Doubler-brush** | **Dumb-brush** |
|---|---|---|---|---|---|---|
| **Diameter (nm)** | 9.7(±1.8) | 16.5(±1.5) | 20(±2.4) | 21.3(±3.0) | 25.5(±2.9) | 20.4(±2.9) |

**Figure S6.** Additional AFM images (top) and average dry-state size measurements (bottom) of PSP bottlebrushes (DP5, DP20 and DP35) and Quasar, Doubler-brush and Dumb-brush.

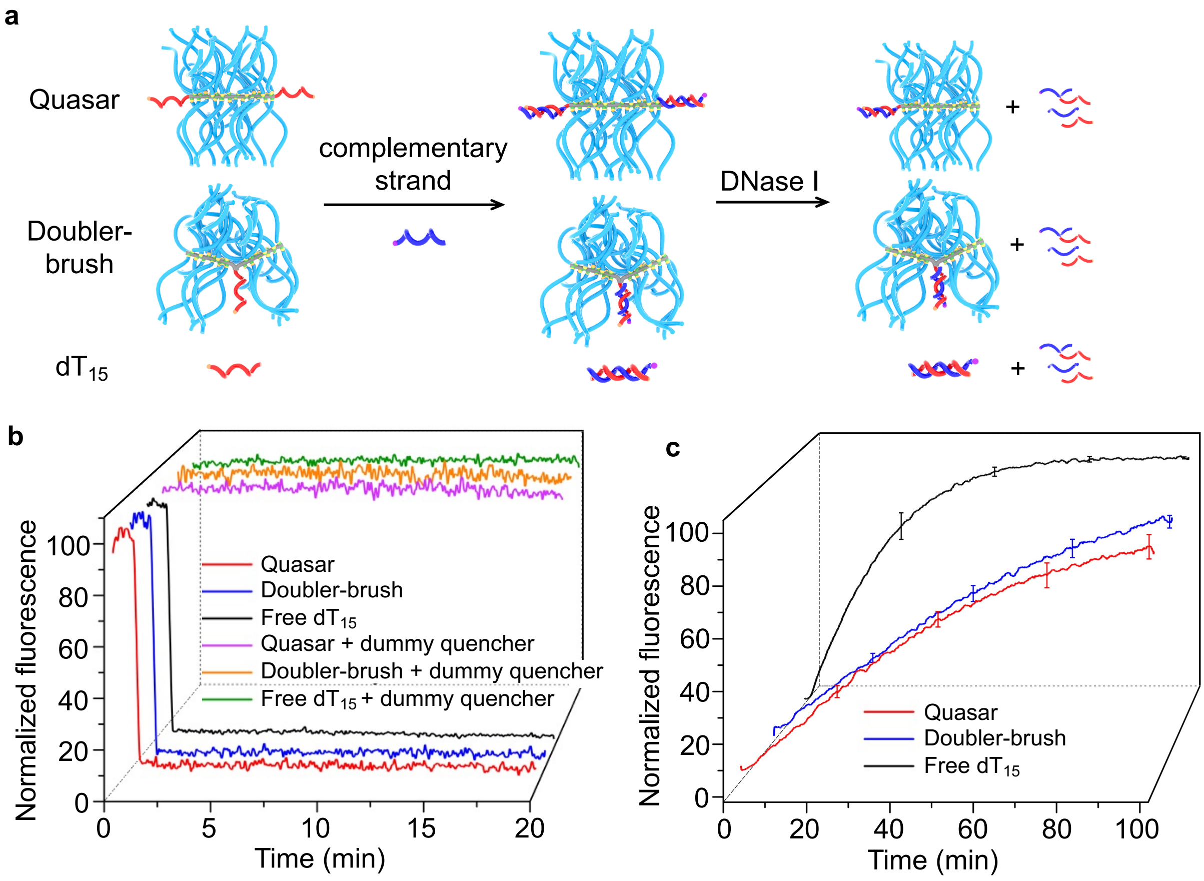


**Figure S7.** Hybridization and enzymatic degradation of $dT_{15}$ Quasars. (a) Schematics of hybridization and enzymatic degradation assay. Hybridization kinetics (b) and DNase I degradation kinetics (c) of $dT_{15}$ Quasars and $dT_{15}$.

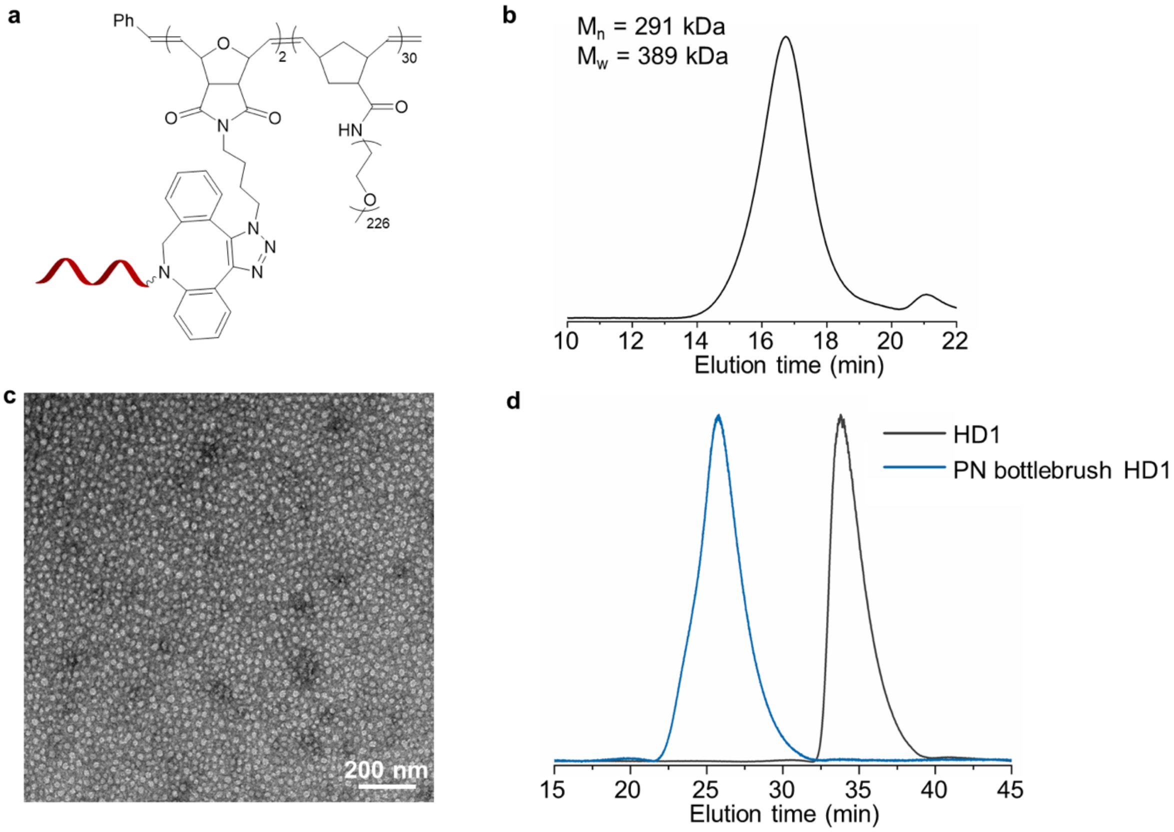


**Figure S8.** Characterizations of PN bottlebrush oligonucleotides conjugates. (a) Structure of PN bottlebrush oligonucleotide conjugates. (b) DMF-GPC chromatogram of PN bottlebrush. (c) Representative TEM images of PN bottlebrush with negative staining (2% uranyl acetate). (d) Aqueous GPC chromatogram of PN bottlebrush HD1 conjugates and free HD1.

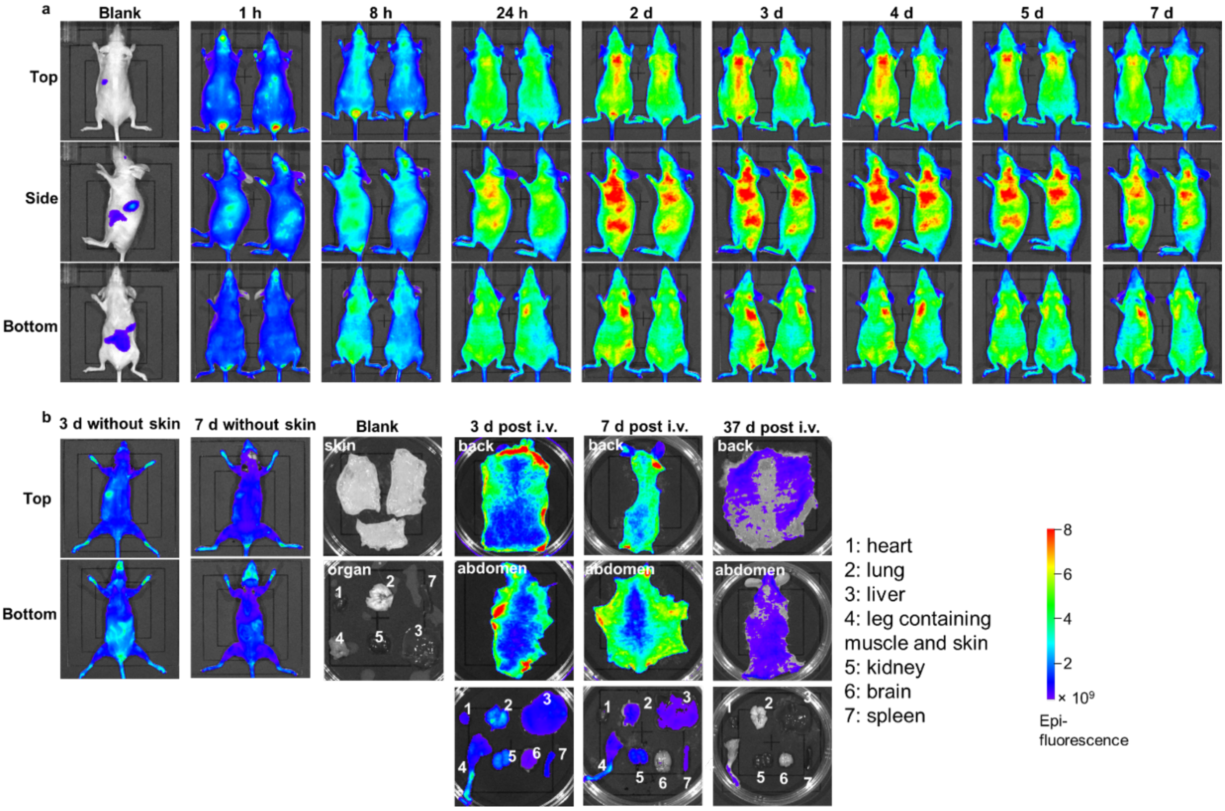


**Figure S9.** Fluorescence monitoring of SKH1-Elite mice dosed intravenously with Cy5-labeled PSP bottlebrush (DP30). (a) Daily imaging of live animals for 7 days. (b) *Ex vivo* imaging of organs 3, 7, and 37 days post injection. Imaging settings were kept identical.

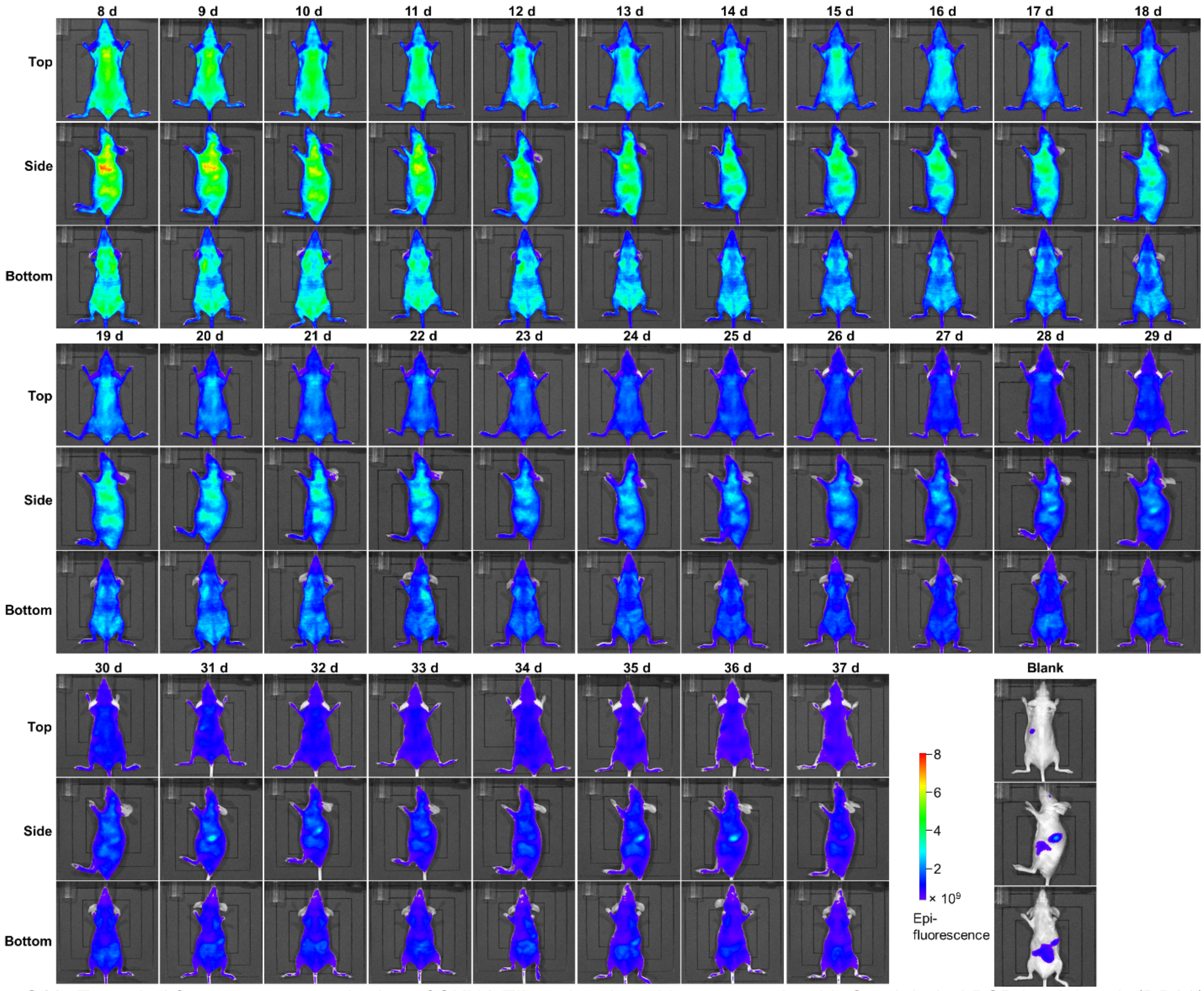


**Figure S10.** Extended fluorescence monitoring of SKH1-Elite mice dosed intravenously with Cy5-labeled PSP bottlebrush (DP30) for 37 days.

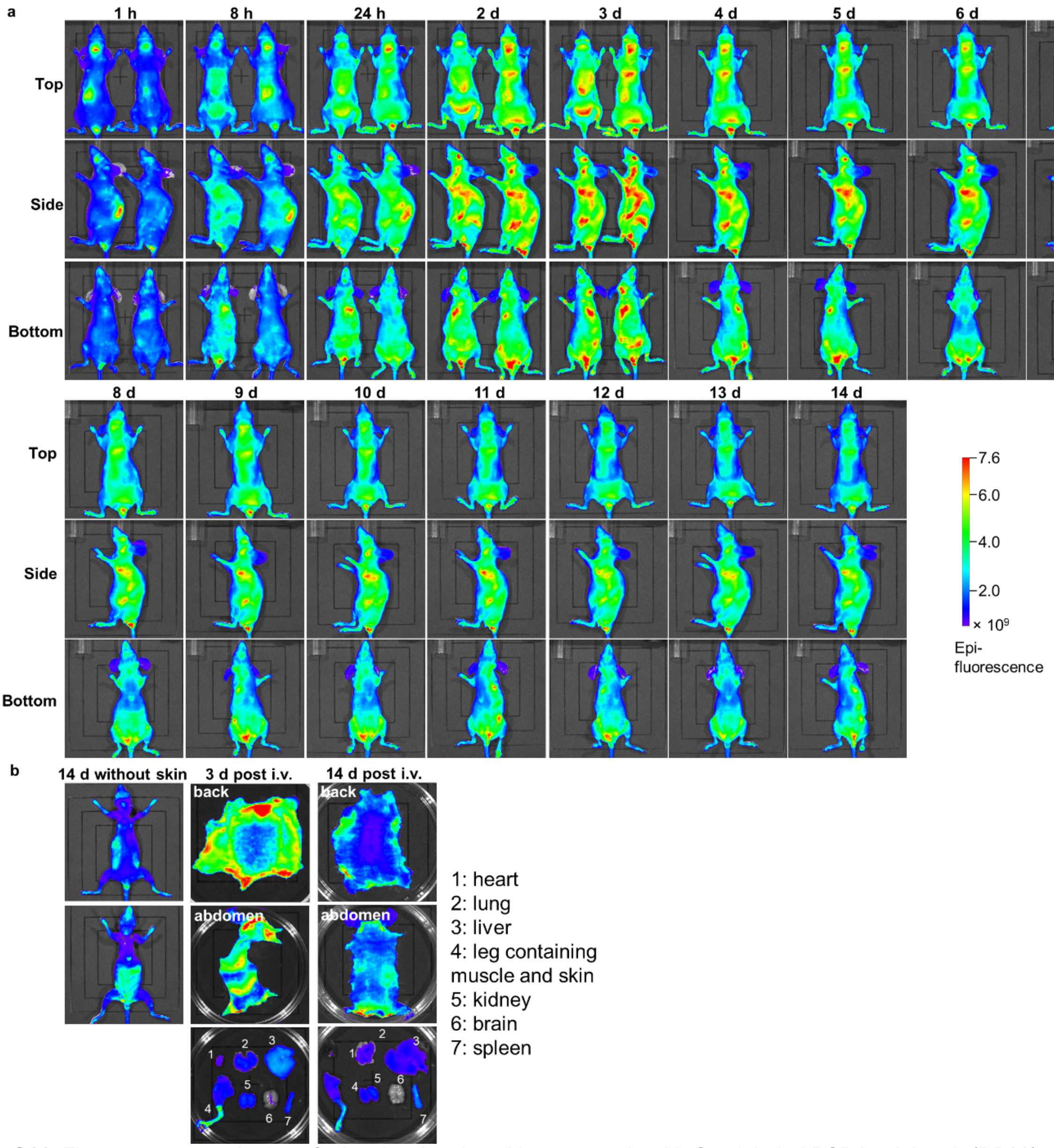


**Figure S11.** Fluorescence monitoring of athymic mice dosed intravenously with Cy5-labeled PSP bottlebrush (DP30). (a) Daily imaging of live animals for 14 days. (b) *Ex vivo* imaging of organs 3 and 14 days post injection. Imaging settings were kept identical.

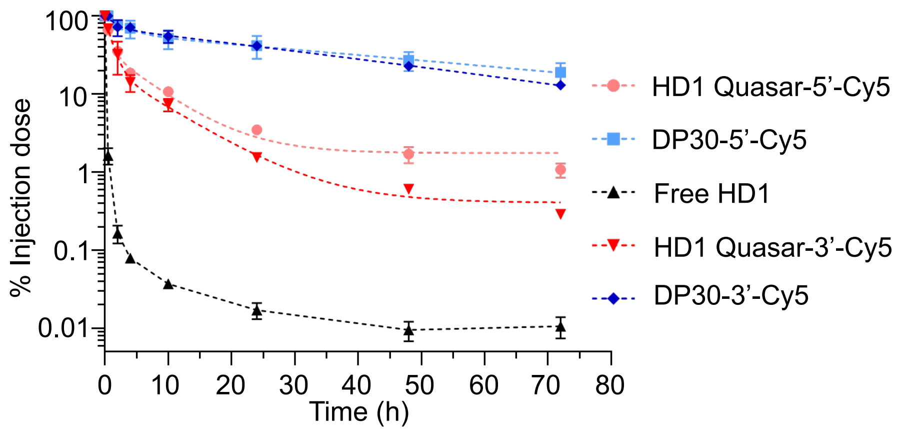


| Sample ID | $t_{1/2\alpha}$ (h) | $t_{1/2\beta}$ (h) | AUC$_\infty$ (nmol/mL·h) |
|---|---|---|---|
| **Free HD1** | 0.25 | 0.43 | 2.12 |
| **DP30-5'-Cy5** | 2.13 | 47.56 | 204.4 |
| **DP30-3'-Cy5** | 1.21 | 31.86 | 188.7 |
| **HD1 Quasar-5'-Cy5** | 0.79 | 5.94 | 34.1 |
| **HD1 Quasar-3'-Cy5** | 0.56 | 4.76 | 24.1 |

**Figure S12.** Plasma pharmacokinetics (top) and calculated parameters (bottom) of 3'- and 5'-Cy5-labeled PSP bottlebrush (DP30-5'-Cy5 and DP30-3'-Cy5) and Quasars in C57BL/6 mice.

### NMR Spectra of compound 1-3 (Scheme S1)

Compound **1**

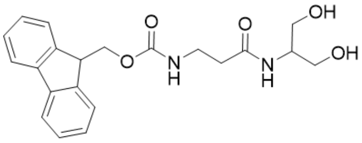

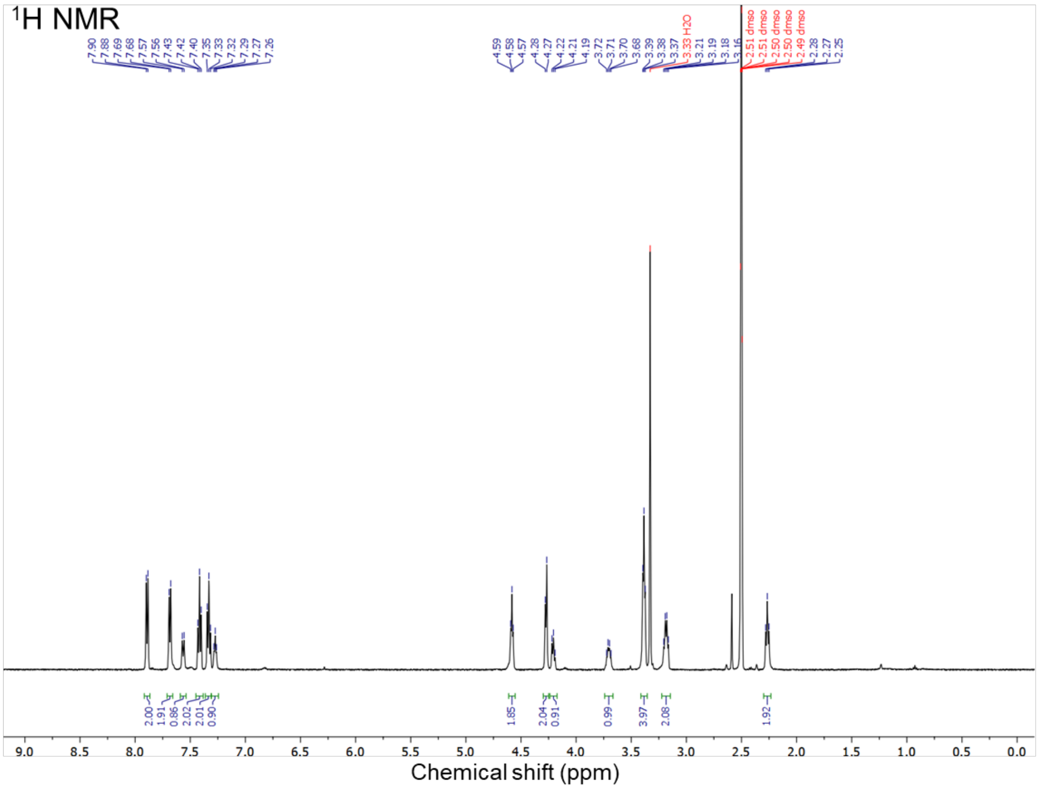


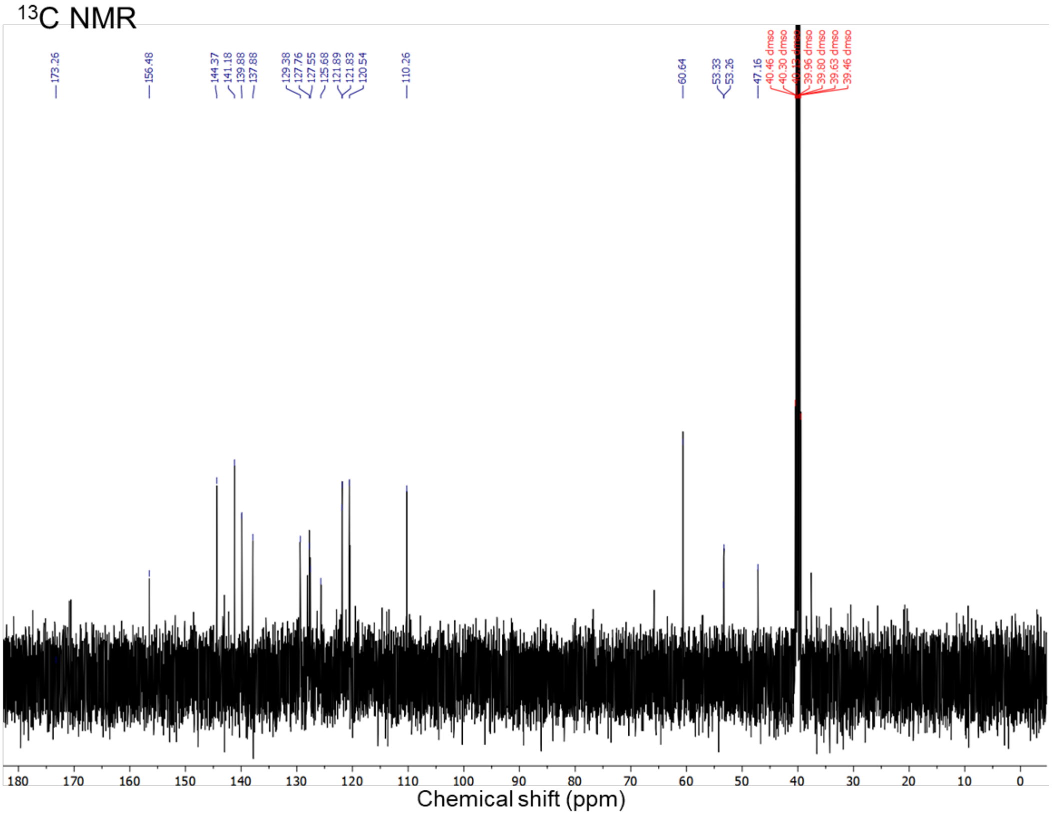

Compound **2**

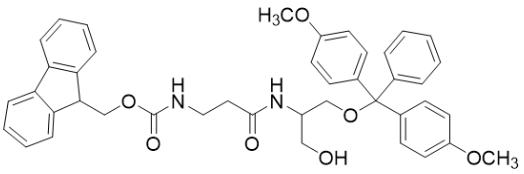


$^1$H NMR

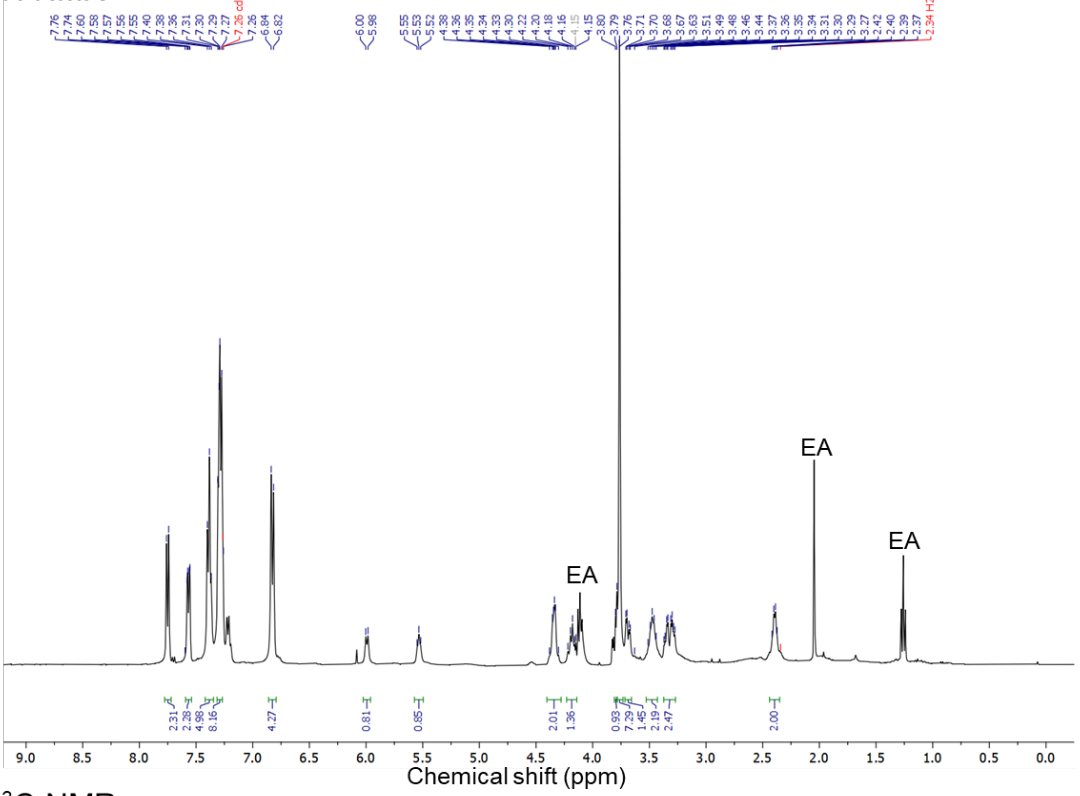


$^{13}$C NMR

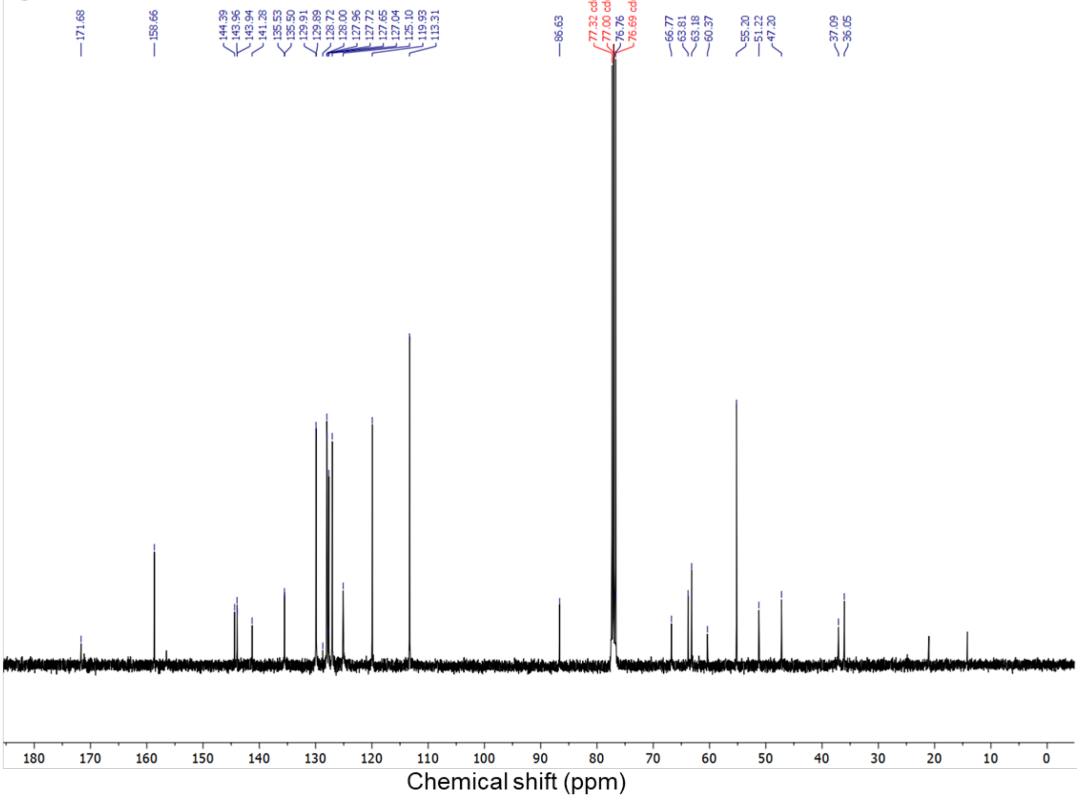

Compound **3**

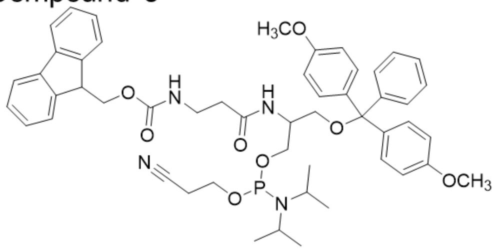


$^1$H NMR

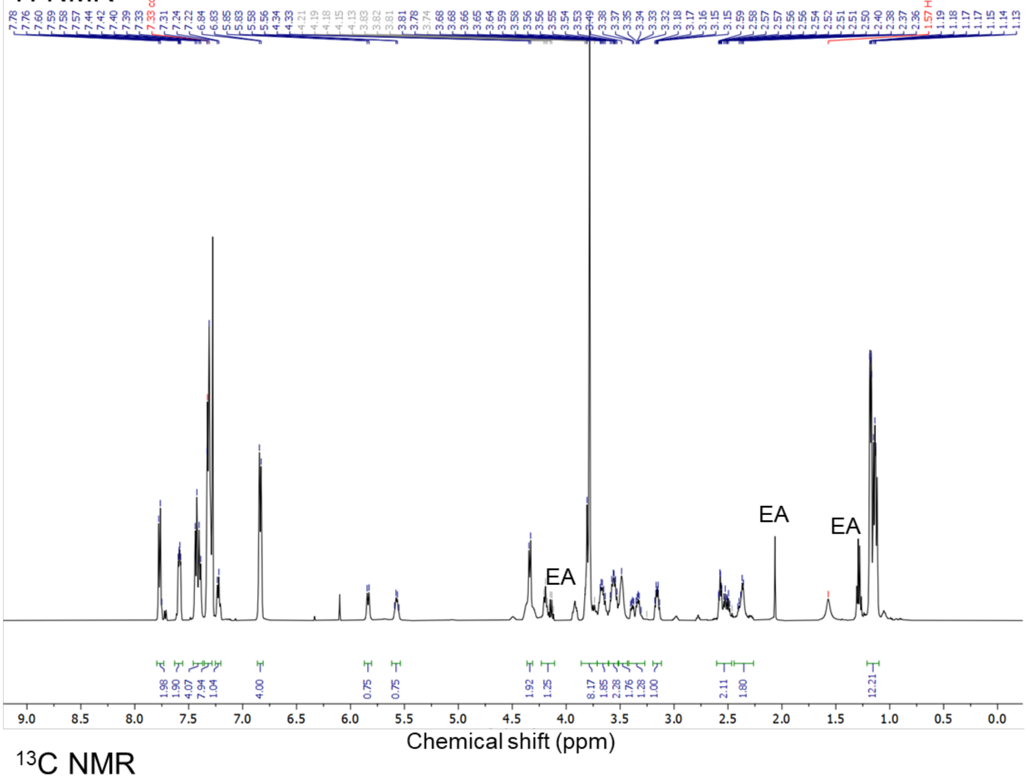


$^{13}$C NMR

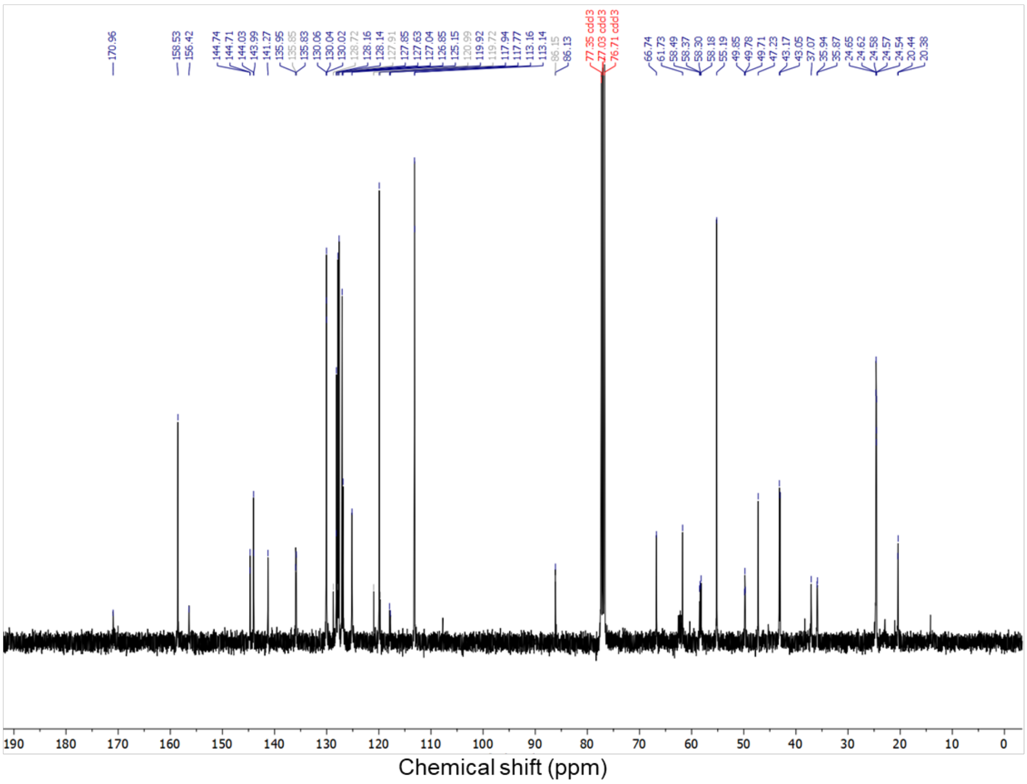


### Author Contributions

K.Z. and Y.W. devised the experiments and wrote the manuscript. Y.W. conducted the synthesis of materials, purification, characterizations, and biological experiments. S.D., Z.Z., and R.W. devised the computational simulations and wrote the relevant sections of the manuscript. Z.Z. and R.W. carried out the simulations. All other authors contributed to material synthesis and/or discussion of the results. All authors edited the manuscript.
